# Non-Canonical Regulation of Phosphatidylserine Metabolism by a Phosphatidylinositol Transfer Protein and a Phosphatidylinositol 4-OH Kinase

**DOI:** 10.1101/696336

**Authors:** Yaxi Wang, Peihua Yuan, Ashutosh Tripathi, Martin Rodriguez, Max Lönnfors, Michal Eisenberg-Bord, Maya Schuldiner, Vytas A. Bankaitis

## Abstract

The phosphatidylserine (PtdSer) decarboxylase Psd2 is proposed to engage in an endoplasmic reticulum (ER)-Golgi/endosome membrane contact site (MCS) that facilitates phosphatidylserine decarboxylation to phosphatidylethanomaine (PtdEtn) in *Saccharomyces cerevisiae*. While this MCS is envisioned to consist of Psd2, the Sec14-like phosphatidylinositol transfer protein (PITP) Sfh4, the Stt4 phosphatidylinositol (PtdIns) 4-OH kinase, the Scs2 tether, and at least one other uncharacterized protein, functional data that address key foundations of this model are sparse. We now report that Psd2, Sfh4 and Stt4 are the only components individually required for biologically sufficient Psd2-dependent PtdEtn production. Surprisingly, neither the PtdIns-transfer activity of Sfh4 nor its capacity to activate Stt4 is required to stimulate the Psd2 pathway. Instead, Sfh4 activates the Psd2 pathway via a specific Sfh4-Psd2 physical interaction. Whereas the data indicate an Sfh4-independent association of Stt4 with Psd2 as well, we find Stt4 also regulates Psd2 activity indirectly by influencing the PtdSer pool accessible to Psd2 for decarboxylation. These collective results demonstrate that the proposed ER-Golgi/endosomal MCS model fails to provide an accurate description of the Psd2 system in yeast, and provide an example where the biological function of a Sec14-like PITP is uncoupled from its ‘canonical’ activity as a PtdIns transfer protein.

## INTRODUCTION

The high degree of biochemical compartmentation of eukaryotic cells affords distinct intracellular organelles the capacity to house unique biological functions. This feature is also an isolating principle that demands cells navigate problems associated with execution of physiologically coherent programs of inter-organellar communication and signaling that are critical for proper cell homeostasis. Whereas the vesicular pathway that connects organelles of the secretory pathway represents one mechanism by which lipids and proteins traffic between compartments, other evidence supports involvements for non-vesicular modes of lipid trafficking as well. That evidence includes early studies of non-vesicular trafficking of PtdSer to mitochondria which were predicated on the assumption that mitochondria represent the sole compartments of residence for the enzyme which decarboxylates PtdSer to PtdEtn. It is now understood that even yeast express two PtdSer decarboxylases– the mitochondrial Psd1 and the endosomal Psd2 (Clancey, Chang et al., 1993, Trotter, Pedretti et al., 1993, Trotter & Voelker, 1995). Both enzymes are posited to utilize membrane contact sites (MCS) for supply of ER-synthesized PtdSer to the compartments of decarboxylation (Voelker, 2005).

MCS are actively maintained foci of close apposition between two distinct organelles and these structures seemingly bridge essentially all cellular compartments (Shai, Yifrach et al., 2018, Valm, Cohen et al., 2017, Wu, Carvalho et al., 2018). Characterization of molecules that populate various MCS complexes defines an intensely active area in contemporary cell biology. In excess of twenty such MCS classes have been identified to date, and specific protein components are assigned to each --including lipid transfer proteins and molecular tethers (Levine & Loewen, 2006, Raychaudhuri & Prinz, 2010, Scorrano, De Matteis et al., 2019, Wu et al., 2018). MCS/lipid trafficking models are conceptually attractive and there is intense speculation regarding how such complexes control lipid flow and distribution between eukaryotic organelles. Direct evidence to that effect remains elusive, however. In many cases, ablation of any single MCS component is without clear biological consequence. Multiple MCS genes need to be ablated before a phenotype is recognized. Further complicating functional analyses, protein tethers often associate with several MCS classes.

Given PtdEtn is an essential lipid in yeast, mutants defective in both Psd1 and Psd2 activities are ethanolamine (Etn) auxotrophs (Figure 1A). This unambiguous biological phenotype offers experimental advantages that recommend the yeast PtdSer decarboxylation system as model for functional dissection of a candidate MCS. The *PSD* system has been exploited to that end and, in the case of Psd2, current models propose that endosomal Psd2-dependent PtdSer decarboxylation requires assembly of an ER-endosomal MCS (Figure 1A) (Riekhof, Wu et al., 2014, Voelker, 2005). Herein, we demonstrate that intrinsic Psd2 activity is independent of the known molecular tethers proposed as components of a Psd2 MCS, or of other proteins that localize to MCS complexes such as the *OSH* proteins. We further show Psd2 interacts with both a PITP and a PtdIns 4-OH kinase independently, and establish the PITP::Psd2 interaction is directly required for biologically-sufficient activation of Psd2. Surprisingly, the activation mechanism, while dependent on the ability of PITP to physically engage Psd2, is independent of PITP activity as a PtdIns-transfer protein. The data also suggest the PtdIns 4-OH kinase involvement on the biological sufficiency of Psd2-mediated PtdSer decarboxylation is complex and is exerted via both direct and indirect mechanisms. The collective data establish that the ER-endosomal MCS model in its present form represents an unsatisfactory description of the Psd2 system in yeast, and demonstrate the need for direct and physiologically relevant assays for mechanistic dissections of MCS function.

**Figure 1.**
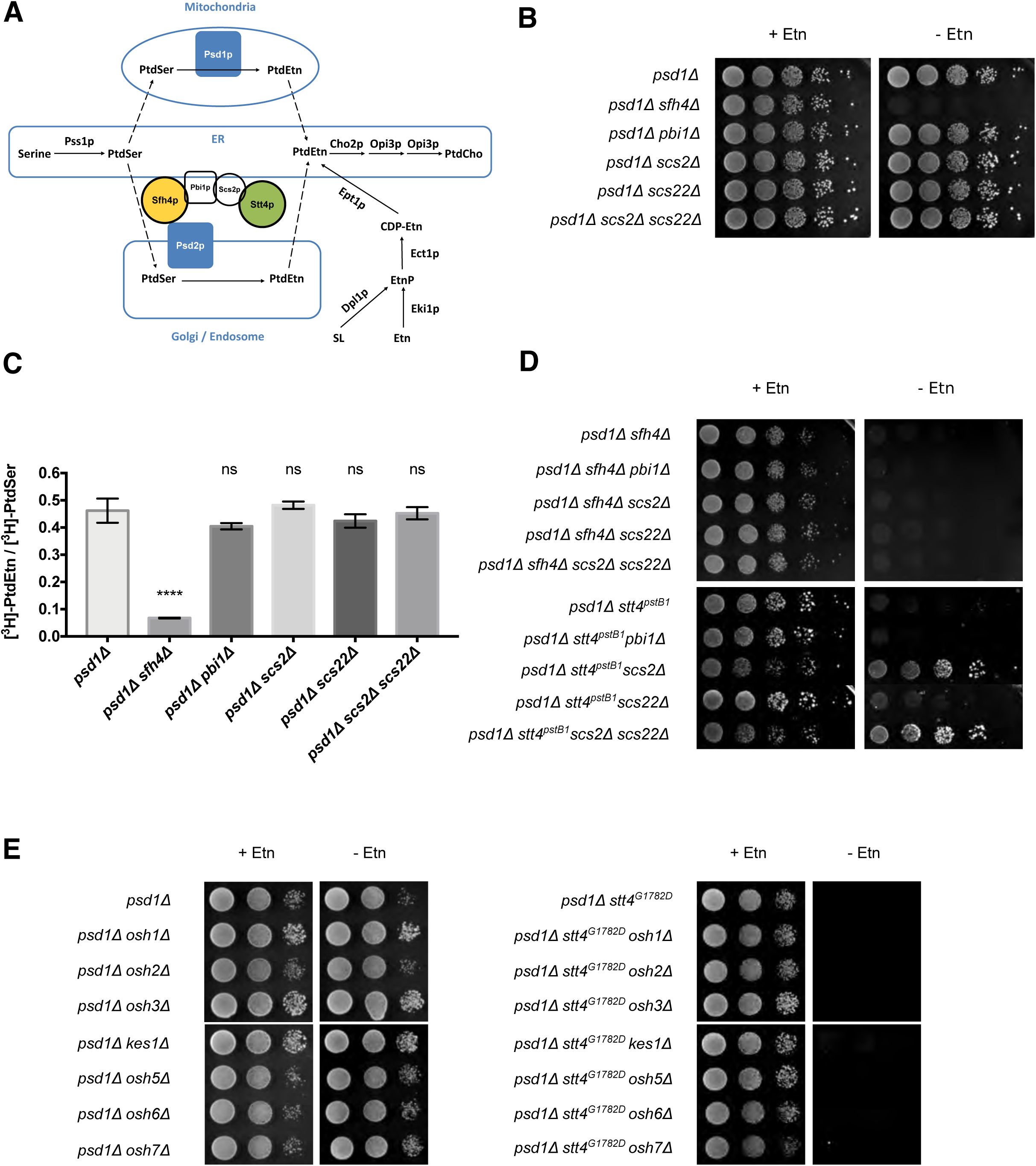
PtdSer decarboxylation and functional characterization of the Psd2 MCS. **(A)** PtdEtn synthesis in yeast occurs via two primary mechanisms. The first involve the decarboxylation of PtdSer catalyzed by either the mitochondial PtdSer decarboxylase Psd1 or the trans-Golgi network/endosomal-localized PtdSer decarboxylase Psd2. Psd2 activity is proposed to require the assembly of an MCS composed of the Sfh4 PITP, the PtdIns-4-OH kinase Stt4, the molecular tether Scs2 that binds Stt4, and the uncharacterized Pbi1. The second major pathway for PtdEtn synthesis involves salvage of exogenous Etn, or Etn-P produced in the terminal step of sphingolipid degradation, via the action of the cytidine diphosphate (CDP)-Etn pathway. **(B)** Yeast strains of the indicated genotype (at left) were dilution spotted onto solid glucose minimal media either supplemented with Etn (2mM) or not as indicated at top. Results were recorded after 48 hours of incubation. Failure to grow in the absence of Etn identifies failure in Psd2 pathway activity as all strains are functionally ablated for Psd1. **(C)** The efficiencies of PtdSer decarboxylation to PtdEtn were measured for *psd1Δ, psd1Δ pbi1Δ, psd1Δ scs2Δ, psd1Δ scs22Δ, psd1Δ scs2Δ scs22Δ* strains by radiolabeling cells with [^3^H]-Serine, extraction of total phospholipid, and quantification of [^3^H]-PtdSer and [^3^H]-PtdEtn (see Materials and Methods). The efficiencies of PtdSer decarboxylation are expressed as [^3^H]-PtdEtn/[^3^H]-PtdSer ratios. Values represent averages from three independent experiments plotted as mean ± standard error of the mean (SEM). Statistical comparisons (ns, p > 0.05; *p ≤ 0.05; **p ≤ 0.01; ***p ≤ 0.001; ****p ≤ 0.0001; in comparison with the *psd1Δ* control) were performed by one-way analysis of variance (ANOVA) with Dunnett multiple comparisons test. **(D)** Yeast strains of the indicated genotype (at left) were dilution spotted onto solid glucose minimal media either supplemented with Etn (2mM) or not as indicated at top. Results were recorded after 48 hours of incubation at 30°C. Restoration of growth in the absence of Etn identifies partial reactivation of the Psd2 pathway in *psd1Δstt4^pstB1^* mutants. **(E)** Isogenic yeast strains with the designated genotypes (at left) were spotted in 10-fold dilution series on solid synthetic complete media with or without Etn (2mM) and incubated at 30°C for 48 hours.

## RESULTS

### VAP-family proteins and Pbi1 are dispensable for Psd2 pathway activity

The *PSD* system has been exploited using various approaches to arrive at a model that posits endosomal Psd2-dependent PtdSer decarboxylation requires assembly of a specialized ER-endosomal contact site composed of Psd2, the Sfh4/Pdr17 PITP, the Stt4 PtdIns 4-OH kinase, the ER-localized VAP-family tethering factor Scs2, and a functionally uncharacterized protein Pbi1 (YPL272C; Figure 1A,B; (Riekhof et al., 2014, Voelker, 2005)). We find no evidence to support positive roles for Scs2 and Pbi1 in Psd2 pathway activity in cells, however. Individual deletion of the *PSD2* and *SFH4* structural genes inactivated PtdSer decarboxylation through the Psd2 pathway -- as assessed by each deletion allele conferring Etn auxotrophy on a *psd1Δ* mutant deficient in the metabolically redundant mitochondrial pathway for PtdSer decarboxylation to PtdEtn (Figure 1B, data not shown). However, individual deletion of either the *PBI1* or *SCS2* structural genes had no such effect. Deletion of the *SCS22* gene, which encodes the second Scs2-like protein in yeast, similarly had no deleterious effect on growth of *psd1Δ* cells on Etn-free media. Indeed, even combinatorial ablation of both the *SCS2* and *SCS22* structural genes failed to impair Psd2-dependent PtdSer decarboxylation in cells as judged by phenotypic criteria (Figure 1B).

We considered the possibility that flux through the Psd2 pathway in cells might operate in excess of biological threshold and that the observed Etn prototrophies failed to report significant, yet phenotypically silent, roles for Scs2 and Pbi1 in promoting metabolic flux through this pathway. That possibility was contraindicated by [^3^H]-serine metabolic labeling experiments that monitored Psd2-dependent conversion of [^3^H]-PtdSer to [^3^H]-PtdEtn in vivo. Using the [^3^H]-PtdEtn/[^3^H]-PtdSer ratio as readout of Psd2 pathway activity, a significant decrease in the [^3^H]-PtdEtn/[^3^H]-PtdSer ratio was measured in the *psd1Δ sfh4Δ* double mutant relative to the *psd1Δ* positive control (Figure 1C). Individual ablation of either Pbi1, Scs2, or Scs22 activities, or combinatorial inactivation of both Scs2 and Scs22, failed to compromise the efficiency of Psd2-mediated PtdSer decarboxylation in cells.

To interrogate potentially antagonistic functional interactions, we tested whether inactivation of Pbi1, Scs2 and/or Scs22 rescued Psd2 pathway activity in *sfh4Δ* and/or *stt4^pstB1^* mutants. The *stt4^pstB1^*allele was identified in the original genetic screen for mutants presumed to be defective in Psd2 activity (Trotter, Wu et al., 1998). In no case did we observe phenotypic rescue upon incorporation of *pbi1Δ*, *scs2Δ* or *scs22Δ* alleles into a *psd1Δ sfh4Δ* genetic background. The isogenic *pbi1Δ*, *scs2Δ* and *scs22Δ* derivatives consistently maintained the Etn auxotrophy characteristic of the parental *psd1Δ sfh4Δ* strain (Figure 1D). Interestingly, *scs2Δ* (but not *pbi1Δ* or *scs22Δ*) enabled significant growth of the *psd1Δstt4^pstB1^*mutant on Etn-free medium. This phenotype was accompanied by a reproducible 1.4-fold increase in [^3^H]-PtdEtn/[^3^H]-PtdSer ratio relative to the isogenic *psd1Δstt4^pstB1^* control (data not shown). Thus, if anything, the VAP-family protein Scs2 interferes with, rather than promotes, Stt4 involvement in Psd2-dependent PtdSer decarboxylation in cells.

### Psd2 pathway activity is not modulated by individual *OSH* protein function

Previous studies established a functional (albeit antagonistic) interaction between a specific member of the yeast oxysterol binding protein family (Kes1 or Osh4) and the major yeast PITP Sec14 in the regulation of PtdIns4*P* signaling (Fang, Kearns et al., 1996, Huang, Mousley et al., 2018, Li, Rivas et al., 2002). Given the proposal that Osh proteins are functional components of intermembrane contact sites (Stefan, Manford et al., 2011, Tian, Ohta et al., 2018), and Osh6 and Osh7 show PtdSer/PtdIns4*P* transfer activity in vitro (Chung, Torta et al., 2015, Moser von Filseck, Copic et al., 2015), we considered the possibility that *OSH* proteins are involved in the Psd2 pathway. However, neither individual deletion of each *OSH* gene, nor *OSH6* or *OSH7* overexpression, was of any consequence to the growth of *psd1Δ* cells on Etn-free medium (Figure 1E, data not shown). In addition, neither deletion of any single *OSH* gene, nor overexpression of *OSH6* or *OSH7*, restored growth to either *sfh4Δpsd1Δ* or *psd1Δstt4^pstB1^*cells on Etn-free medium (Figure 1E, data not shown). Psd2 pathway activity was similarly indifferent to functional ablation of the structural genes encoding other tethers (i.e. the newly recognized endosomal tether Vps13 and ER-Golgi tethers Lam5 or Lam6; (De, Oleskie et al., 2017, Elbaz-Alon, Eisenberg-Bord et al., 2015, Gatta, Wong et al., 2015, Kumar, Leonzino et al., 2018, Murley, Sarsam et al., 2015)) when assessed phenotypically in *psd1Δ* genetic backgrounds (data not shown). Importantly, while all of these directed assays could have overlooked an important factor not previously suggested in the literature, a systematic survey of all non-essential yeast genes failed to identify any functions other than Sfh4 and Psd2 that are required for growth of *psd1Δ* cells in Etn-free medium (Suppl. Figure S1, Suppl. Table S1). These cumulative analyses support the idea that Sfh4 and Stt4 are the two primary components that individually exert significant control over Psd2-mediated PtdSer decarboxylation in cells.

### Stt4 PtdIns 4-OH kinase activity is required for Etn prototrophy in cells expressing Psd2 as sole PtdSer decarboxylase

Several lines of experimentation were conducted to assess whether Stt4 enzymatic activity was required for the Etn prototrophy of cells expressing Psd2 as sole source of PtdSer decarboxylase activity. As first step, the *stt4^pstB1^* allele was recovered from genomic DNA and the *STT4* nucleotide sequence determined in its entirety. *STT4* encodes a 1901 amino acid PtdIns 4-OH kinase, and the *stt4^pstB1^* allele differed from the wild-type *STT4* nucleotide sequence by a single G→A transition mutation located within the Stt4 catalytic domain (Figure 2A). This domain is bounded by residues 1641 and 1848 [http://pfam.xfam.org/protein/P37297], and the *pstB1* mutation substitutes Gly with Asp at amino acid 1782 (Figure 2A). We henceforth refer to the *stt4^pstB1^* allele as *stt4^G1782D^*.

**Figure 2.**
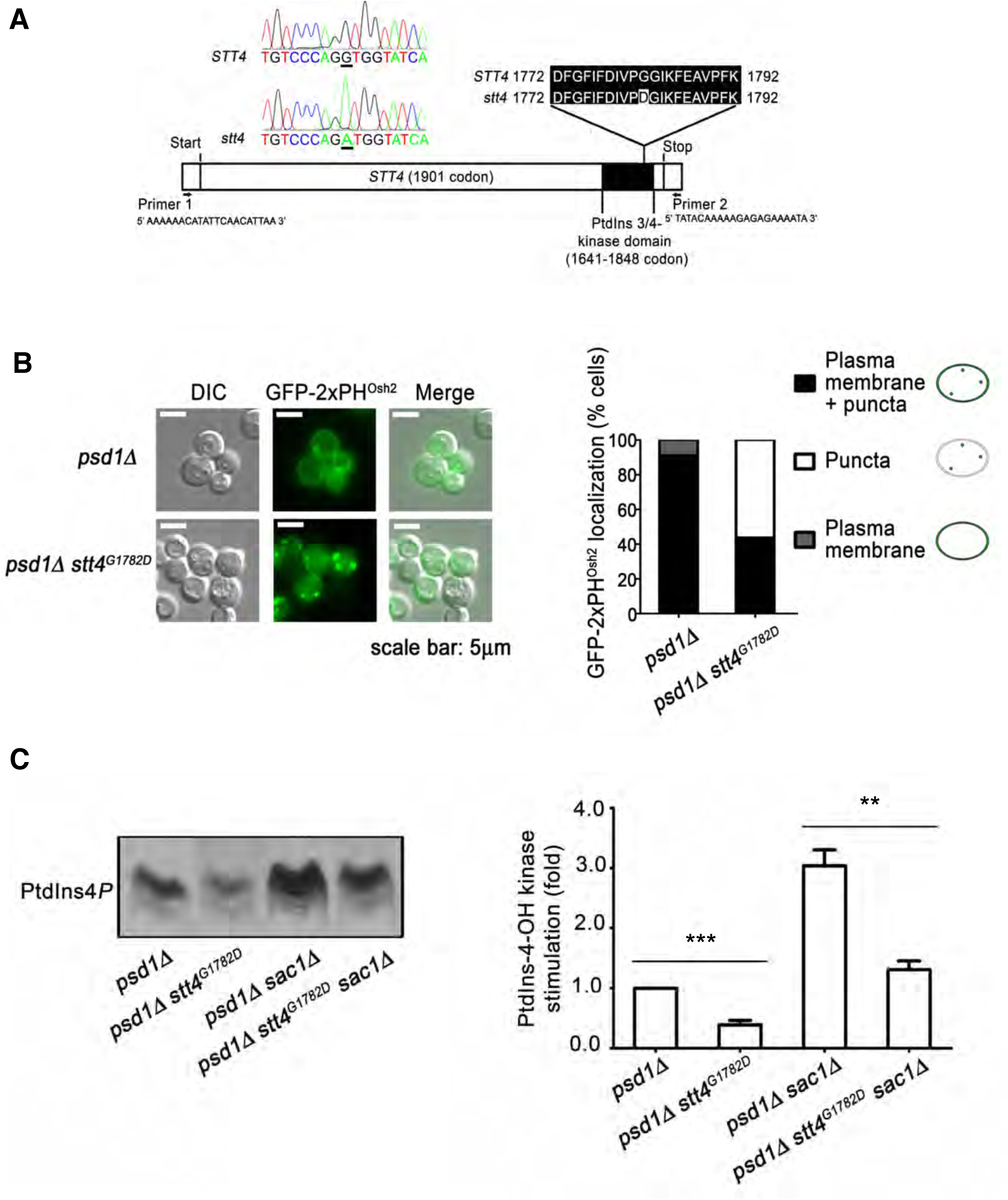
Stt4 PtdIns 4-OH kinase activity is required for biologically sufficient Psd2 activit*y*. **(A)** The full-length *STT4* gene was recovered from the *pstB1* mutant genome and DNA sequence analysis identifies a G_1782_D missense substitution in the PtdIns-kinase domain. **(B)** Distributions of the GFP-2xPH^Osh2^ PtdIns4*P* biosensor in *psd1Δ* and *psd1Δ stt4^G1782D^* cells were determined by fluorescence microscopy imaging and classified for each cell imaged as plasma membrane localization, punctate TGN/endosomal localization, or both (left panel). The fractional representation of cells exhibiting each distribution profile is shown (right panel). The plasma membrane localization of the biosensor reads out the Stt4-dependent PtdIns4*P* pool. **(C)** Appropriate yeast strains (genotypes at bottom) were radiolabeled to steady-state with [^3^H]-inositol and shifted to 37°C for 3 hr. The *sac1Δ* strains were used to enhance the PtdIns4*P* signal and to emphasize the PtdIns4*P* synthesis contribution to the data. Phospholipids were extracted, resolved by thin layer chromatography and autoradiography (left panel). PtdIns4*P* band intensities were measured by densitometry (arbitrary units) and expressed as a ratio of PtdIns4*P* to total input c.p.m (input range from 1070088 to 1627675 c.p.m). The normalized ratios were related to a *psd1Δ* control (set to 1.0; right panel). Values represent averages from three independent experiments plotted as mean ± SEM. Statistical comparisons of values used the ‘unpaired two-tailed t-test”. ***p ≤ 0.001 in comparison to the *psd1Δ* strain; **p ≤ 0.01 in comparison to the *psd1Δ sac1Δ* strain.

Two independent assays were employed to interrogate whether Stt4^G1782D^ is intrinsically defective in PtdIns4*P* production in situ. First, GFP-2xPH^Osh2^ was used as a PtdIns4*P* biosensor in vital imaging assays (Roy & Levine, 2004). This biosensor registers an Stt4-dependent plasma membrane PtdIns4*P* pool as well as a punctate TGN/endosomal PtdIns4*P* pool generated by the Pik1 PtdIns 4-OH kinase (Nile, Tripathi et al., 2014). Some 91% of the *psd1Δ STT4* control cells displayed localization of the biosensor to both plasma membrane and punctate TGN/endosomal structures. A minor fraction of cells exhibited a solely plasma membrane localization profile (Figure 2B). By contrast, only 44% of *psd1Δ stt4^G1782D^* cells showed detectable plasma membrane staining with this reporter (along with punctate TGN/endosomal staining), and 56% of cells displayed solely punctate TGN/endosomal localization of the reporter with no visible plasma membrane staining at all (Figure 2B). Those imaging results reported compromise of Stt4-mediated PtdIns4*P* production in *stt4^G1782D^*cells.

Supporting data were obtained from metabolic [^3^H]-inositol radiolabeling experiments coupled to analyses of Stt4-dependent PtdIns4*P* production. These measurements exploited *sac1Δ* mutants deficient in the Sac1 PtdIns4*P* phosphatase – i.e. the dominant activity for degradation of Stt4-generated PtdIns4*P* in yeast (Foti, Audhya et al., 2001, Guo, Stolz et al., 1999, Nemoto, Kearns et al., 2000, Rivas, Kearns et al., 1999). As a result, these measurements directly reported Stt4 PtdIns 4-OH kinase activity in vivo. Indeed, *stt4^G1782D^* mutants displayed significant reductions in PtdIns4*P* levels relative to isogenic *STT4* control cells (Figure 2C).

To directly confirm that PtdIns 4-OH kinase activity was required for Stt4 function in promoting Psd2-mediated PtdSer decarboxylation in cells, the consequences of expressing a catalytic dead version of Stt4 in *psd1Δ stt4^G1782D^* yeast were determined. Design of the catalytic-dead *stt4^D1754A^*allele was guided by previous characterization of the kinase-dead *pik1^D918A^*allele (Strahl, Hama et al., 2005). Low copy yeast centromeric *(CEN)* plasmids carrying *STT4, stt4^G1782D^ or stt4^D1754A^* expression cassettes driven by the natural *STT4* promoter were used in these analyses. Whereas individual expression of *STT4* or *stt4^G1782D^*was sufficient to rescue viability of an *stt4^ts^* mutant strain at a restrictive temperature of 37°C, *stt4^D1754A^*expression failed to do so (Suppl. Figure S2A). When these same expression constructs were introduced into the *psd1Δ stt4^G1782D^* mutant, only *STT4* expression supported growth of *psd1Δ stt4^G1782D^*cells on Etn-free medium (Suppl. Figure S2B). Thus, both *stt4^G1782D^* and *stt4^D1754A^* scored as defective alleles. These data indicated that: (i) PtdIns 4-OH kinase activity of Stt4 was required for its involvement in the Psd2 pathway, (ii) Stt4^G1728D^ represents a functional hypomorph with reduced in vivo lipid kinase activity, and (iii) this crippled enzyme, while sufficiently functional to support yeast cell viability, was inadequate for restoration of Etn prototrophy to yeast expressing Psd2 as sole PtdSer decarboxylase activity.

The yeast Stt4 PtdIns 4-OH kinase catalytic domain requires accessory proteins for biologically sufficient activity and it assembles into two distinct complexes in that regard. Complex I consists of the Stt4 catalytic subunit with the associated Ypp1 and Erf3 subunits, whereas complex II consists of Stt4 and the accessory subunits Ypp1 and Sfk1 (Baird, Stefan et al., 2008). Of these, Sfk1 is the only activity dispensible for cell viability. We found that *psd1Δ sfk1Δ* double mutant yeast were not compromised for growth on Etn-free media (Suppl. Figure S2C), indicating that Stt4 complex II was not required for Psd2 pathway activity in vivo.

Overexpression of Ypp1, Efr3 or Sfk1 neither evoked Etn auxotrophy in *psd1Δ* yeast, nor did enhanced expression of these non-catalytic subunits rescue the Etn auxotrophy of *psd1Δ stt4^G1782D^* cells (Suppl. Figure S2D).

### The PtdIns-binding substructure of Sec14-like PITPs is conserved in Sfh4

Sec14 and its related PITPs potentiate PtdIns 4-OH kinase activities in cells (including Stt4) by a presentation mechanism that renders PtdIns a superior substrate for the lipid kinase (Bankaitis, Mousley et al., 2010, Schaaf, Ortlund et al., 2008). The biological specificities that come with the presentation model are striking as, of the six Sec14-like PITPs in yeast, Sfh4 is unique in its involvement in potentiating metabolic flux through the Psd2 pathway (Suppl. Figure S3A). Thus, we considered the possibility that Sfh4 acts in a canonical fashion by enhancing Stt4 activity to expand production of a PtdIns4*P* pool dedicated to activation of the Psd2 pathway. This hypothesis predicted that Sfh4 mutants unable to bind PtdIns would be defective in the context of Psd2-mediates PtdSer decarboxylation. To identify Sfh4 residues critical for PtdIns-binding, the Sfh4 primary sequence was threaded onto a high-resolution crystal structure of Sfh3 bound to PtdIns. Sfh3 is the most similar of the yeast PITPs to Sfh4 (49% primary sequence identity and 65% similarity). The optimized Sfh4::PtdIns homology model described a Sec14-fold comprised of eleven α-helices, six 3_10-_helices, and five β-strands (Suppl. Figure S3B) (Schaaf et al., 2008, Sha, Phillips et al., 1998). The common PtdIns-coordination substructure essential for specific PtdIns binding observed in crystal structures of other Sec14-like proteins was also clearly recognized in the Sfh4 structural model (Suppl. Figure S3C) (Phillips, Sha et al., 1999, Ren, Pei-Chen Lin et al., 2014, Schaaf et al., 2008). Of note in that regard are Sec14 residues T_236_ and K_239_ which play critical roles in PtdIns headgroup binding and are conserved throughout the Sec14 superfamily (Schaaf et al., 2008). The Sfh4 cognates of those residues are Sfh4^T266^ and Sfh4^K269^ (Suppl. Figure S3D), and Sfh4^T266D,K269A^ and Sfh4^T266W,K269A^ double mutants were expected to be defective for PtdIns-binding based on electrostatic and steric considerations.

### Biochemical validation of Sfh4 mutants defective in PtdIns-binding

Consistent with expectations, both Sfh4^T266D,K269A^ and Sfh4^T266W,K269A^ exhibited strong PtdIns-binding deficits as determined by three independent assays. First, recombinant Sfh4, Sfh4^T266D,K269A^ and Sfh4^T266WK269A^ proteins were purified from *E.coli* and queried for PtdIns-transfer activity in vitro using an assay that quantifies mobilization of [^3^H]-PtdIns from donor rat liver microsomes to acceptor PtdCho vesicles. Whereas both Sfh4 and the Sec14 positive controls displayed robust PtdIns-transfer, neither Sfh4^T266D,K269A^ nor Sfh4^T266W,K269A^ exhibited significant activity (Figure 3A). Second, as independent confirmation of the [^3^H]-PtdIns transfer assay, the activities of Sfh4 and mutant proteins were assayed in a real-time fluorescence de-quenching PtdIns-transfer system (Somerharju, van Loon et al., 1987). In this assay, transfer is scored by increased fluorescence of pyrene-labeled PtdIns ([Pyr]-PtdIns) upon its mobilization from the quenching environment provided by the donor vesicles. Again, Sfh4 catalyzed robust transfer of [Pyr]-PtdIns, whereas Sfh4^T266D,K269A^ was strongly deficient (Suppl. Figure S3E). Third, whereas enhanced *SFH4* expression rescued growth of a *sec14-1^ts^* mutant at the restrictive temperature of 37°C, neither elevated expression of Sfh4^T266D,K269A^ nor of Sfh4^T266W,K269A^ elicited detectable phenotypic rescue of the *sec14-1^ts^*growth defect (Figure 3B). This failure was not the trivial consequence of mutant Sfh4 protein instability as immunoblotting experiments demonstrated both mutant proteins accumulated in cells to levels comparable to those of Sfh4 itself (Figure 3C).

**Figure 3.**
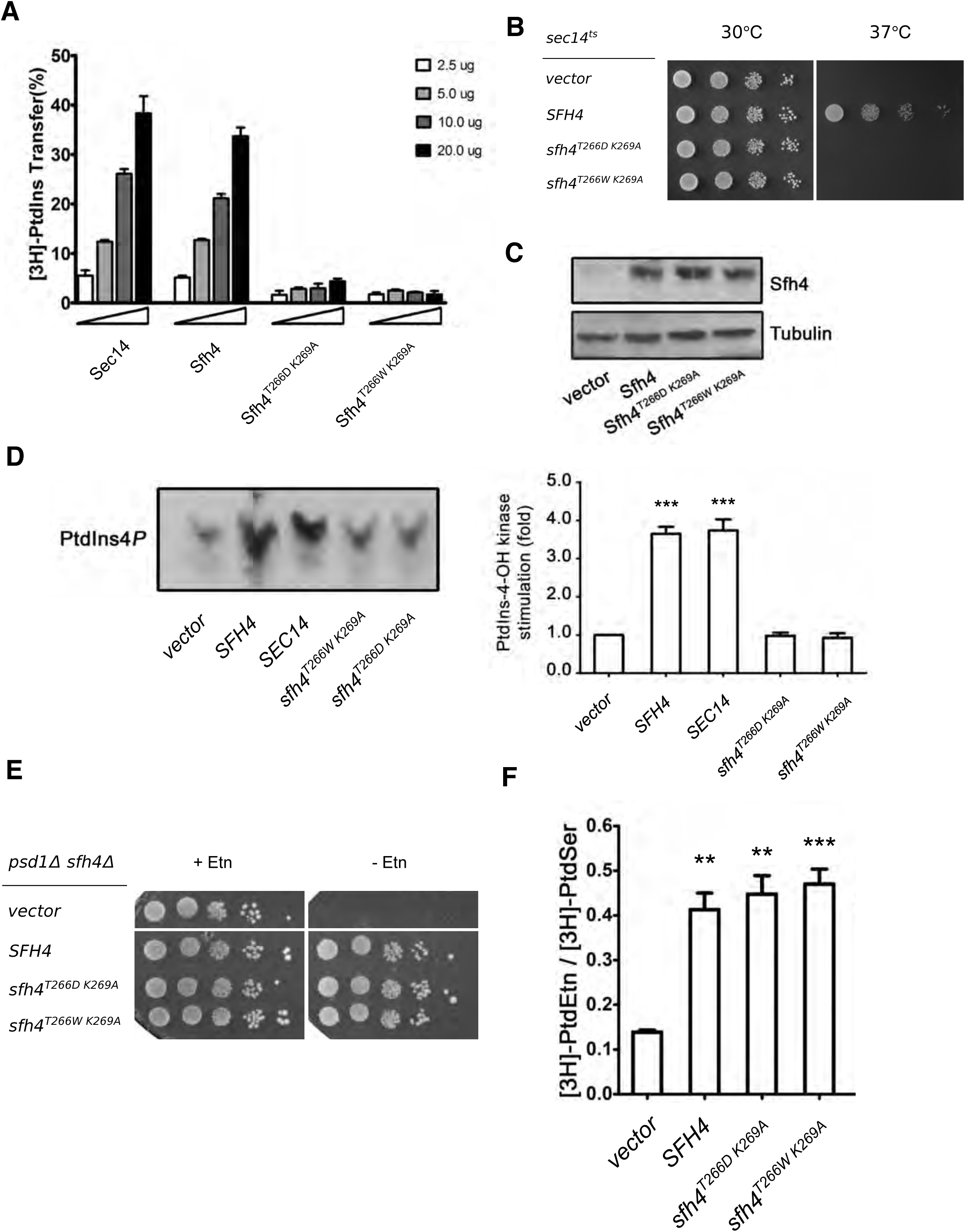
Sfh4 mutants defective for PtdIns-binding retain ability to potentiate Psd2 activity in vivo. **(A)** PtdIns transfer assays for wild-type and mutant Sfh4 proteins. The indicated step series of purified recombinant Sec14, Sfh4, Sfh4^T266D^ ^K269A^ and Sfh4^T266W^ ^K269A^ were assayed for PtdIns-transfer from rat liver microsomes to acceptor vesicles. Transfer activity is expressed as the ratio of [^3^H]-PtdIns in the acceptor vesicle fraction to total input [^3^H]-PtdIns in donor vesicle fraction X 100%. Values represent averages from three independent experiments plotted as mean ± SEM. [^3^H]-PtdIns inputs ranged in these assays from 5559-5935 cpm, with background transfer ranging from 546 –670 cpm (determined in mock samples where buffer was incorporated rather than protein). **(B)** Isogenic *sec14^ts^* yeast transformed with a mock YCp vector or YCp vectors carrying *SFH4* WT, *sfh4^T266D^ ^K269A^* or *sfh4^T269W^ ^K269A^* expression cassettes under *PGK1* promoter control were spotted in 10-fold dilution series on uracil-free media and incubated at the permissive- and restrictive temperatures of 30°C and 37°C respectively. Growth at 37°C signifies the protein of interest is competent for PtdIns binding/exchange. **(C)** Immunoblots of lysates prepared from *psd1Δ sfh4Δ* yeast carrying the designated YCp vector where expression of the proteins of interest was driven by the strong constitutive *PGK1* promoter. Equal amounts of total protein were loaded. Sfh4 and tubulin (normalization control) were visualized with anti-Sfh4 or anti-tubulin antibodies, respectively. **(D)** Left panel: Isogenic *sec14^ts^ sac1 sfh4Δ* yeast mutants ectopically expressing the indicated Sec14 or Sfh4 proteins from a yeast centromeric vector where the expression cassette was driven by the *PGK1* promoter were radiolabeled to steady-state with [^3^H]-inositol and shifted to 37°C for 3 hours. Phospholipids were extracted, and PtdIns4*P* resolved by thin layer chromatography ((Phillips et al., 1999, Schaaf et al., 2008); see Materials and Methods). The vector and *SEC14* derivatives represented negative and positive controls, respectively. Right panel: PtdIns4*P* band intensities were measured by densitometry and expressed as a ratio of PtdIns4*P/*total input counts (inputs ranged from 1332937 to 1804138 c.p.m). These normalized ratios were related to a vector control (set to 1.0 on the relative scale). Values represent averages from three independent experiments plotted as mean ± SEM. Statistical comparisons of values used the ‘unpaired two-tailed t-test”. ***p ≤ 0.001 in comparison to a vector control. **(E)** Isogenic *psd1Δ sfh4Δ* yeast transformed with mock YCp vectors or YCp vectors expressing the indicated genes whose expression was driven by the *SFH4* promoter, were spotted in 10-fold dilution series on uracil-free media with or without supplementation with 2mM Etn as indicated at top. Plates were incubated at 30°C for 48 hours. **(F)** Psd2-mediated PtdSer decarboxylation is supported by Sfh4 mutants defective in PtdIns binding/exchange. The yeast strains described for panel E above were incubated with [^3^H]-serine for 3 hours at 30°C, phospholipids were extracted, and PtdSer and PtdEtn resolved by TLC. Values represent the mean ± SEM from at least 3 independent experiments. Statistical comparisons of values used the ‘unpaired two-tailed t-test” relative to vector control (**p ≤ 0.01, ***p ≤ 0.001).

Finally, [^3^H]-inositol radiolabeling experiments confirmed that both Sfh4^T266D,K269A^ and Sfh4^T266W,K269A^ were defective in stimulating Stt4-mediated PtdIns4*P* production in vivo. Those measurements were conducted in a *sac1Δ sec14-1^ts^*double mutant background using a previously established experimental design (Ile, Kassen et al., 2010, Schaaf et al., 2008). Briefly, the *sac1Δ* allele inactivates the primary pathway for degradation of the PtdIns4*P* produced by Stt4. The *sec14-1^ts^* allele erases the dominant stimulatory effect of Sec14 on Stt4 in vivo activity when cells are incubated at 37°C -- thereby providing an assay for assessing the stimulatory effects of enhanced expression of Sfh4 and mutants of interest on Stt4 activity. Whereas *SFH4* expression supported a ca. 2.5-fold stimulation of PtdIns4*P* synthesis relative to vector control, cells expressing Sfh4^T266D,K269A^ or Sfh4^T266W,K269A^ exhibited only basal PtdIns4*P* levels that were comparable to those of the negative control (Figure 3D). Thus, while Sfh4 effectively stimulated Stt4 activity in vivo, neither Sfh4^T266D,K269A^ nor Sfh4^T266W,K269A^ were able to do so.

### Sfh4 mutants defective in PtdIns-binding maintain competence in promoting Psd2 pathway activity in vivo

Contrary to expectations that Sfh4 must bind PtdIns to function in the Psd2 pathway, expression of either Sfh4^T266D,K269A^ or of Sfh4^T266W,K269A^ was sufficient to restore Etn prototrophy to a *psd1Δ sfh4Δ* double mutant yeast strain. This level of rescue was phenotypically indistinguishable from that observed for the Sfh4 control even when the mutant proteins were expressed from cassettes driven by the weak *SFH4* promoter (Figure 3E). This result deserves special emphasis as the expression cassettes were all carried on low copy centromeric vectors. That is, a configuration that supported a physiologically appropriate level of Sfh4 expression. Further testimony to the functionality of these PtdIns-binding defective Sfh4 mutants was provided by [^3^H]-serine labeling experiments. The efficiencies of Psd2-dependent conversion of [^3^H]-PtdSer to [^3^H]-PtdEtn were comparable irrespective of whether Sfh4, Sfh4^T266D,K269A^ or Sfh4^T266W,K269A^ represented the sole sources of Sfh4 activity in the cell (Figure 3F).

That activation of the Psd2 pathway was not a simple result of Sfh4-stimulated PtdIns4*P* production by the Stt4 PtdIns 4-OH kinase was independently indicated by two other lines of evidence. First, inactivation of the Sac1 PtdIns4*P* phosphatase was insufficient to effect either a ‘bypass Sfh4’ condition for Psd2-dependent decarboxylation of PtdSer in *psd1Δ sfh4Δ* mutants, or to rescue Psd2 pathway activity in *psd1Δ stt4^G1782D^* mutants (Suppl. Figure S4A). Genetic ablation of Sac1 PtdIns4*P* phosphatase activity elevates the levels of PtdIns4*P* generated by Stt4 some 10-fold (Audhya & Emr, 2002, Cleves, Novick et al., 1989, Foti et al., 2001, Guo et al., 1999, Nemoto et al., 2000, Rivas et al., 1999). Metabolic labeling data confirmed that *sac1Δ* failed to improve the efficiency of Psd2-dependent conversion of [^3^H]-PtdSer to [^3^H]-PtdEtn in either mutant background (data not shown). Those negative results were also recapitulated in genetic backgrounds where each of the other yeast phosphoinositide phosphatase structural genes were individually deleted (Suppl. Figure S4B).

### Isolation of Sfh4 mutants specifically defective in stimulating Psd2 activity

To determine what specific Sfh4 properties are required for its positive involvement in Psd2-dependent PtdSer decarboxylation, we sought Sfh4 mutants with defects restricted to the Psd2 activation function. To that end, an unbiased genetic screen was designed to identify Sfh4 mutants specifically defective in Psd2 activation. This screen coupled mutagenic PCR with gap-repair of a centromeric *SFH4* expression plasmid in a *ura3 sfh4Δ psd1Δ sec14^ts^*recipient strain (Suppl. Figure S5A). Expression of a mutant Sfh4 incompetent for activation of the Psd2 pathway would fail to impart Etn prototrophy to the strain. The recipient strain had the additional feature that it could not grow at 37°C because it carries a *sec14-1^ts^*allele. This phenotypic feature was exploited to filter away trivial *sfh4* loss-of-function mutations as the repaired plasmid drives expression of the mutagenized *SFH4* open reading frame from the powerful constitutive *PGK* promoter -- a configuration sufficient to support growth of the *sec14-1^ts^*recipient at 37°C if the expressed Sfh4 protein is a functional PITP. Thus, gap-repaired Ura^+^ transformants of the *ura3 sfh4Δ psd1Δ sec14^ts^*recipient strain were selected and screened by replica plating for colonies displaying the dual unselected traits of: (i) Etn auxotrophy (mutant Sfh4 fails to activate the Psd2 pathway), and (ii) the ability to form colonies at 37°C (mutant Sfh4 retains sufficient PITP activity to act as functional surrogate for Sec14).

From a total of >8,000 Ura^+^ transformants so analyzed, a single isolate was identified that exhibited the desired unselected phenotypes of Etn auxotrophy and ability to grow at 37°C. The plasmid was recovered from the yeast isolate and re-transformed into a naive *ura3 sfh4Δ psd1Δ sec14^ts^*recipient strain using Ura^+^ selection. The Ura^+^ transformants again displayed the unselected Etn auxotrophy and thermoresistant growth traits -- thereby confirming plasmid linkage of these hallmark phenotypes. Nucleotide sequence analyses of the plasmid-borne *SFH4* identified a single TTT→CTT transition mutation in *SFH4* codon 175 that substituted Phe at that position with Leu. Mapping of this substitution onto the threaded Sfh4 structural model projected that residue Phe175 lies on the surface of the open conformer of this PITP at a position far removed from the substructure that coordinates PtdIns headgroup binding (Figure 4A).

**Figure 4.**
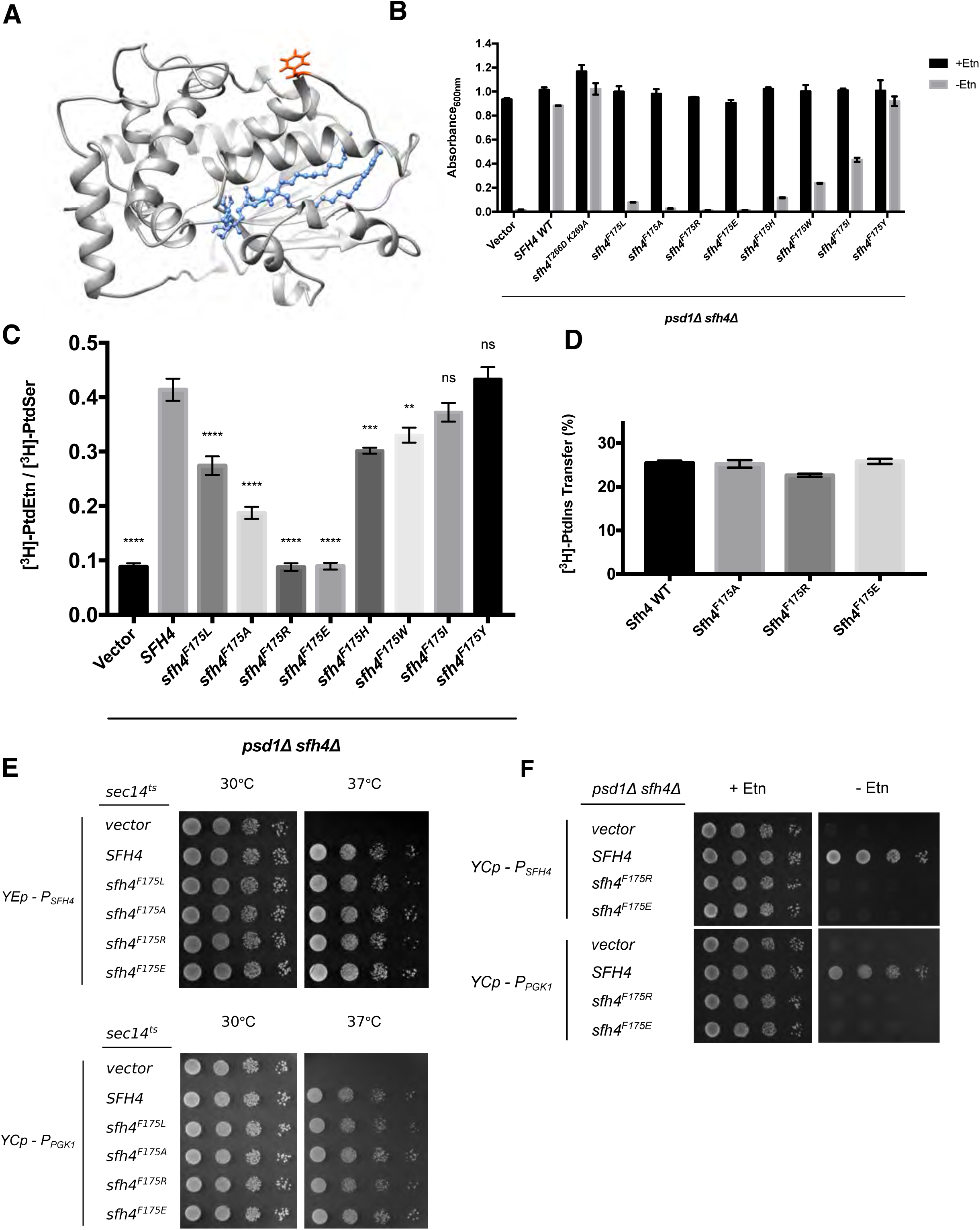
Sfh4 mutants specifically defective in Psd2 pathway stimulation. **(A)** Sfh4 homology model based on threading onto the Sfh3 crystal structure is shown and residue F_175_ is highlighted in orange-red. The bound PtdIns molecule is rendered in blue ball and stick mode. **(B)** Growth phenotypes in Etn (2mM)-containing (black bars) or Etn-free (gray bars) minimal medium lacking uracil of a parental *psd1Δ sfh4Δ* strain transformed with mock YCp(*URA3*) vector or YCp (*URA3*) vectors expressing either *SFH4* or individual Sfh4^F175^ missense mutants (indicated at bottom). Cultures were initially grown in uracil-free Etn-replete minimal SD medium at 30°C to mid-logarithmic phase. Cells were washed twice with water, and re-suspended in uracil-free Etn-replete minimal SD medium. After a 4 hour incubation at 30°C to deplete any residual Etn stores, the cultures were washed one more time with water and diluted to an A_600_ of 0.01 in the appropriate minimal SD medium. The diluted cultures were incubated at 30°C for 46.5 hours, and the A_600_ of each culture was recorded. Values represent averages from two independent experiments (n=2) plotted as mean ± range. **(C)** PtdSer flux through the Psd2 pathway was measured and is plotted as described above in the legends to Figures 1C. Ectopic expression of the Sfh4 control and the test Sfh4^F175^ variants in the *psd1Δ sfh4Δ* parental strain was in all cases driven by an *SFH4* promoter cassette carried on a low copy *CEN* plasmid. Values represent averages from at least three independent experiments plotted as mean ± SEM. Statistical comparisons (ns, p > 0.05; *p ≤ 0.05; **p ≤ 0.01; ***p ≤ 0.001; ****p ≤ 0.0001; in comparison with the *SFH4* control) were performed by one-way analysis of variance (ANOVA) with Dunnett multiple comparisons test. **(D)** Purified recombinant Sfh4, Sfh4^F175A^, Sfh4^F175R^ and Sfh4^F175E^ proteins (10μg) were assayed for [^3^H]-PtdIns-transfer activity as described in the Methods. Transfer activities are expressed as ratios of [^3^H]-PtdIns in acceptor vesicle fractions relative to total input [^3^H]-PtdIns present in donor vesicle fractions X 100%. Values represent averages from three independent experiments plotted as mean ± SEM. [^3^H]-PtdIns inputs in these assays ranged from 11317–11724 cpm, while backgrounds ranged from 342 –394 cpm **(E)** Isogenic *sec14^ts^*yeast transformed with an empty YEp vector or YEp vectors driving *SFH4* WT or *sfh4* mutant genes expression under *SFH4* promoter control (top panels), and isogenic *sec14^ts^* yeast transformed with empty YCp vector or YCp vectors driving ectopic expression of *SFH4* or *sfh4* mutant gene expression from the *PGK1* promoter (bottom panels) were spotted in 10-fold dilution series on uracil-free media and incubated at the permissive- and restrictive temperatures of 30°C and 37°C respectively. Growth at 37°C signifies ability of the corresponding protein to bind/exchange PtdIns and stimulate PtdIns 4-OH kinase activity in vivo. **(F)** Parental *psd1Δ sfh4Δ* yeast were transformed with a YCp(*URA3*) vector or derivatives harboring expression cassettes where transcription of *SFH4* or mutant *sfh4* genes was driven from either the *SFH4* promoter (*P_SFH4_)* or the powerful *PGK1* promoter (*P_PGK1_*). Transformants were spotted in 10-fold dilution series on uracil-free media with or without exogenous Etn (2mM) as indicated at top. Plates were incubated at 30°C for 48 hours.

### Characterization of F_175_ substitutions reveals a gradient of functional defects

To probe the involvement of F_175_ in Sfh4 engagement with the Psd2 pathway, this residue was altered to a series of other amino acids of differing size, hydrophobicity and charge. The abilities of the corresponding Sfh4 mutant proteins to activate the Psd2 pathway were then assessed. In this paradigm, expression of the mutant proteins of interest was driven by the natural *SFH4* promoter and the expression cassettes were introduced into the *psd1Δ sfh4Δ* parent strain on a centromeric low copy (CEN) vector to approximate physiological levels of protein expression. The readout for Etn prototrophy was conducted in liquid medium as this assay is a stricter gauge of Psd2 pathway activity than is the plate assay.

Sfh4^F175Y^ supported growth of the *psd1Δ sfh4Δ* strain in Etn-free media as efficiently as did expression of Sfh4 or expression of the PtdIns-binding-defective Sfh4^T266D,K269A^ mutants (Figure 4B). Sfh4^F175W^ and Sfh4^F175H^ were compromised in their ability to support cell growth in the absence of Etn, but this defect was not as severe as that recorded for Sfh4^F175L^. The growth phenotypes were buttressed by [^3^H]-serine metabolic radiolabeling assays where the [^3^H]-PtdEtn/[^3^H]-PtdSer ratio for Sfh4^F175Y^-expressing cells was essentially the same as that of Sfh4-expressing cells. By contrast, the [^3^H]-PtdEtn/[^3^H]-PtdSer ratios of Sfh4^F175W^ and Sfh4^F175H^ cells were lower than those of Sfh4-expressing cells, but higher than those of cells reconstituted with Sfh4^F175L^ (Figure 4C). Sfh4^F175A^ –expressing cells were even more defective in stimulating Psd2 activity than were cells where Sfh4^F175L^ was the sole source of Sfh4 function (Figure 4B,C). The most extreme defects were scored in cells relying on Sfh4^F175R^ and Sfh4^F175E^ for stimulation of Psd2 activity. Both mutant proteins scored as completely defective (Figure 4B,C).

For all Sfh4^F175^ mutants, the functional deficiencies in stimulating Psd2 pathway activity in vivo were on display even though Sfh4^F175L^, Sfh4^F175A^, Sfh4^F175R^, and Sfh4^F175E^ exhibited essentially wild-type [^3^H]-PtdIns transfer activities in vitro (Figure 4D, data not shown). All of these mutant proteins similarly displayed wild-type rates of [Pyr]-PtdIns transfer in real-time fluorescence de-quenching assays (Suppl. Figure S5B). Moreover, Sfh4^F175R^ and Sfh4^F175E^ expression driven by either the weak *SFH4* promoter from a high-copy episomal plasmid-borne cassette, or driven by the powerful *PGK1* promoter from a low-copy *CEN* plasmid, was sufficient to rescue growth of *sec14^ts^* yeast at the restrictive temperature of 37°C (Figure 4E). Yet, ectopic Sfh4^F175R^ and Sfh4^F175E^ expression from low-copy *CEN* plasmids, where transcription was driven by either the weak *SFH4* promoter or the powerful *PGK1* promoter, failed to restore growth of those cells in Etn-free media. These data confirmed that both mutant proteins were specifically and dramatically defective for Psd2 pathway activation (Figure 4F).

### Sfh4^F175^ substitutions compromise physical interaction with Psd2 in vivo

Previous studies claim Sfh4 binds Psd2, although no study addressed the physiological relevance of such an interaction (Gulshan, Shahi et al., 2010, Riekhof et al., 2014). The availability of mutant Sfh4 proteins that show undiminished PITP activity, yet are strongly defective in stimulation of Psd2 pathway activity in vivo, afforded a system with which to directly interrogate whether Sfh4 and Psd2 engaged in a functionally relevant interaction. To this end, a co-precipitation assay was developed where a C-terminal HA-tagged Psd2 was expressed from a multicopy plasmid in a yeast strain carrying a TAP-tagged *SFH4* allele transplaced into its natural genomic locus.

Indeed, Sfh4 and Psd2 exhibited an interaction as scored by efficient recovery of the tagged C-terminal 17kDa Psd2 α-subunit in an Sfh4-TAP co-precipitation regime (Figure 5A). We conclude this interaction occurred in situ, and was not the trivial artifact of some post-lysis association, as the co-precipitation was not reproduced when cells expressing only Sfh4-TAP were mixed with equal numbers of cells expressing only C-terminally HA-tagged Psd2 and the mixed suspension was lysed in the same vessel. Moreover, this interaction was judged to be specific as no Psd2 was detected in control co-precipitations that offered a TAP-tagged version of the closely related Sfh3 PITP as bait (Figure 5A).

**Figure 5.**
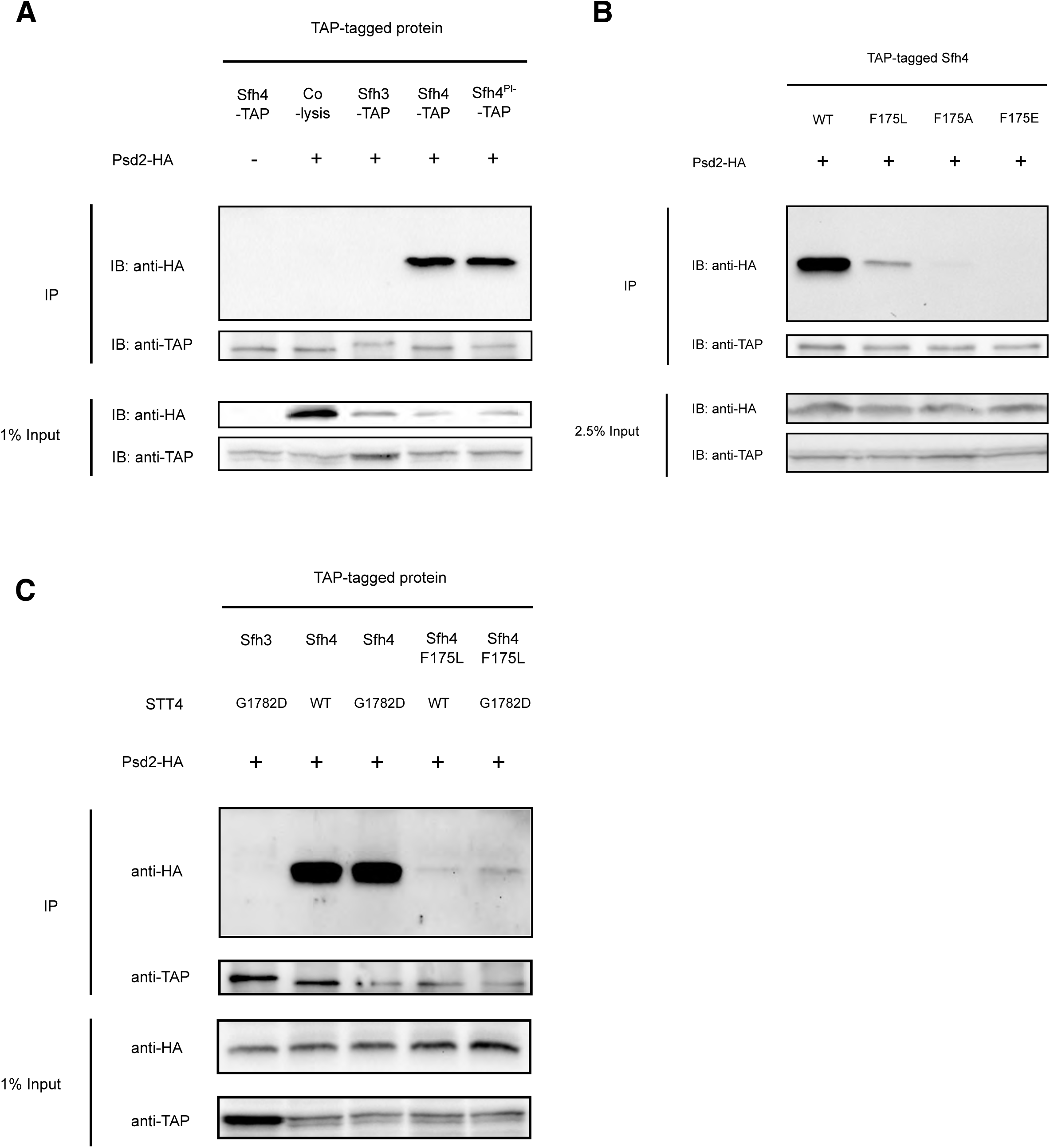
Sfh4 interacts with Psd2 in vivo. **(A)**: Wild-type yeast cells with integrated *SFH3-TAP, SFH4-TAP* or *sfh4^PI-^-TAP* (*sfh4^T266D,K269A^-TAP*) cassette at their endogenous genome loci, were transformed with yeast episomal plasmids driving Psd2-HA expression from the *PSD2* promoter (+) or with a mock episomal vector (-). Transformants were cultured to mid-logarithmic growth phase, collected, mechanically disrupted and clarified lysates produced. Cell lysate aliquots (12 mg total protein) were subjected to anti-TAP immunoprecipitation (IP). All immunoprecipitated samples and input samples were processed for SDS-PAGE and immunoblotting (IB) with anti-HA or anti-TAP antibodies. In the “co-lysis” sample, WT yeast harboring YEp(*PSD2-HA*) and *SFH4-TAP* expressing yeast were grown separately, equal amounts (as measured by A_600_) of each culture were mixed together and lysed in the same tube. The immunoprecipitation procedure was carried out as described above. **(B)** Wild-type yeast cells with integrated *SFH4-TAP, sfh4^F175L^-TAP, sfh4^F175A^-TAP & sfh4^F175E^-TAP* cassettes at their endogenous genomic locus were tested for their abilities to co-precipitate Psd2-HA expressed from a YEp vector as described in (A). Yeast cells carrying either a WT *STT4* locus or the mutant *stt4^G1782D^* allele, and with integrated *SFH4-TAP, sfh4^F175L^-TAP, sfh4^F175A^-TAP & sfh4^F175E^-TAP* cassettes at their endogenous genomic *SFH4* locus, were tested for their abilities to co-precipitate Psd2-HA expressed from a YEp vector as described in (A). The *stt4^G1782D^* was transplaced into the endogenous *STT4* genomic locus via allele exchange.

To determine whether Sfh4^F175^ mutants were compromised for the Psd2 interaction, the co-precipitation assay was run with TAP-tagged Sfh4^F175L^, Sfh4^F175A^ and Sfh4 ^F175E^ mutants. Those representative mutants span a range of deficiencies in Psd2 activity in vivo (in terms of magnitude of severity; Sfh4^F175E^ > Sfh4^F175A^ > Sfh4^F175L^). The corresponding missense substitutions were incorporated into the genomic *SFH4-TAP* allele, the mutant Sfh4^F175^-TAP proteins were recovered by affinity purification from cell-free lysates, and the co-precipitation fractions were interrogated for the HA-tagged Psd2 α-subunit. Satisfyingly, affinity purification of Sfh4^F175L^-TAP demonstrated a strong diminution in capture of the Psd2 α-subunit, whereas Sfh4^F175A^-TAP was even more defective in this assay and Sfh4 ^F175E^-TAP failed to capture any Psd2 α-subunit at all (Figure 5B). Thus, the severities of Psd2-interaction defects for the Sfh4^F175^ mutants recapitulated proportionately the severities of the defects in Psd2-pathway activity in cells expressing these Sfh4^F175^ mutants as sole sources of Sfh4. Importantly, immunoblotting experiments demonstrated that, in all cases, the input cell-free lysate fractions contained similar levels of the HA-tagged Psd2 α-subunit. Those results indicated that neither Psd2 expression/stability, nor the auto-processing activity of this enzyme, was compromised in cells expressing Sfh4^F175^ mutants as their sole source of Sfh4.

### Stt4 deficiencies do not compromise Sfh4-Psd2 interaction

To determine whether the in vivo Psd2::Sfh4 interaction was Stt4-dependent, we tested whether complex formation was compromised in the *stt4^G1782D^* genetic background. To that end, the *stt4^G1782D^* allele was transplaced into the *SFH4-TAP* yeast strain, clarified lysates were generated, and the co-precipitation assay repeated. As shown in Figure 5C, neither the Psd2-Sfh4 nor the crippled Psd2::Sfh4^F175L^ interactions were compromised in *stt4^G1782D^*cells – even though the associated Stt4 deficiencies were of sufficient magnitude to render activity of the Psd2 pathway biologically insufficient for PtdEtn production. The fact that Stt4 deficiencies did not further diminish the already inefficient Psd2::Sfh4^F175L^ interaction argued strongly that Stt4 catalytic activity had no significant role in chaperoning Psd2-Sfh4 complex formation.

### Stt4 interacts with Psd2 in an Sfh4-independent manner

Analogous co-precipitation strategies using Stt4-TAP as bait were conducted to assess potential Stt4-Psd2 interactions (Figure 6). Indeed, both Stt4-TAP and Stt4^G1782D^-TAP were able to capture Psd2-HA. This interaction was considered specific on the basis of three independent controls: (i) significant co-precipitation was not evident in co-lysis experiments controlling for post-lysis association of Stt4-TAP and Psd2-HA, (ii) no Psd2-HA was recovered in co-precipitations using a TAP-tagged Pik1 PtdIns 4-OH kinase as bait (thereby reproducing the *in vivo* functional specificity where the Stt4 PtdIns 4-OH kinase functions in Psd2 pathway activation whereas the Pik1 PtdIns 4-OH kinase does not; data not shown), and (iii) Psd2-HA was not captured by Sfh3-TAP (Suppl. Figure S6A). A caveat of note regarding these co-precipitation experiments is that trace levels of post-lysis Stt4-TAP::Psd2-HA association were detected in co-lysis controls. But, as the levels of Psd2 co-precipitation with Stt4 were much lower in co-lysis controls relative to the test, our interpretation of the data is that the results indicate a genuine Stt4::Psd2 interaction in vivo.

**Figure 6.**
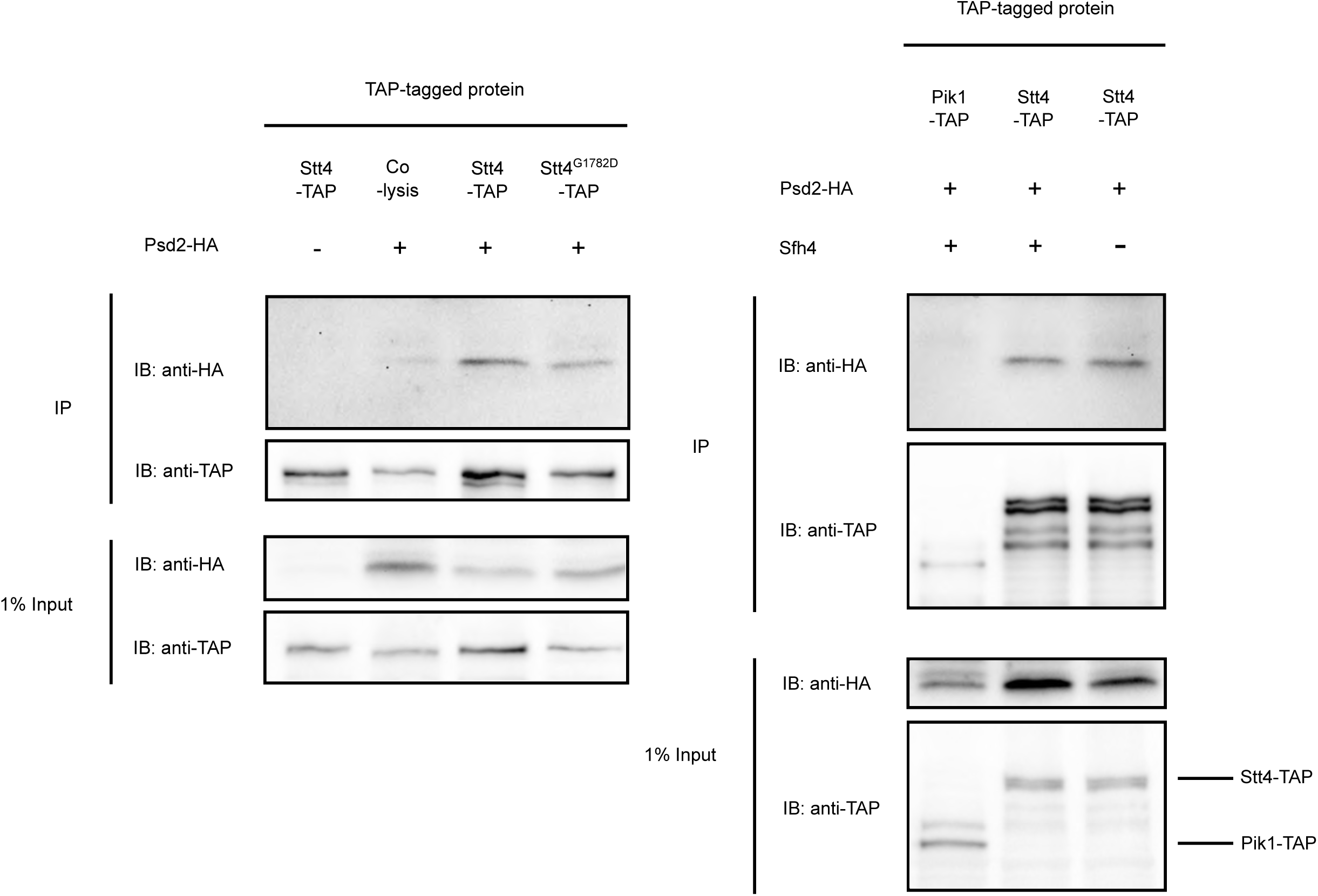
Stt4 interaction with Psd2. Wild-type or *sfh4Δ* yeast carrying integrated *STT4-TAP*, *stt4^G1782D^-TAP* or *PIK1-TAP* cassettes at their endogenous genomic loci (as indicated) were transformed with either a YEp(*PSD2-HA*) (+) or mock YEp vector (-), and the TAP-precipitation regimen was carried out as described in Figure 5. The “co-lysis” control is directly analogous to that described in Figure 5A and scores post-lysis interaction of Psd2-HA and Stt4-TAP.

Interestingly, the Stt4-Psd2 interaction was preserved not only when Stt4^G1782D^-TAP was used as bait, but also when co-precipitations were conducted in the context of an *sfh4Δ* genetic background (Figure 6). Those collective data indicate that Stt4 not only interacted with Psd2 in an Sfh4-independent manner, but that a biologically defective Stt4 impaired for catalytic activity remained competent for this interaction. No physical interaction was scored between Stt4-TAP and Sfh4-HA regardless of whether the co-precipitation experiment was conducted in WT or in *stt4^G1782D^* genetic backgrounds (Suppl. Figure S6B).

### PtdIns4*P* homeostasis indirectly modulates Psd2-dependent PtdSer decarboxylation

During the course of this study, we made several paradoxical observations that indicated a more complex involvement of Stt4-dependent PtdIns4*P* metabolism in PtdSer decarboxylation by Psd2. The first indication came when we observed that *psd1Δ sac1Δ* double mutants, but not *psd2Δ sac1Δ* double mutants, were Etn auxotrophs. That is, inactivation of the Sac1 phosphatase that degrades the PtdIns4-P pool produced by Stt4 paradoxically phenocopied the effects of reduced Stt4^G1782D^ PtdIns 4-OH kinase activity in Psd1-deficient cells (Suppl. Figure S7). What Sac1- and Stt4-deficient cells do share in common is that both exhibit significantly reduced fractional contributions of PtdSer to total cellular phospholipid mass (Rivas et al., 1999, Tani & Kuge, 2014). To assess whether *stt4^G1782D^*similarly influences bulk cellular PtdSer levels, cells were metabolically radiolabeled with [^3^H]-serine, lipids resolved by thin-layer chromatography, and PtdSer was collected and quantified by scintillation counting. The incorporation values obtained were normalized by culture A_600_ and expressed as a relative value to WT control (Figure 7A).

**Figure 7.**
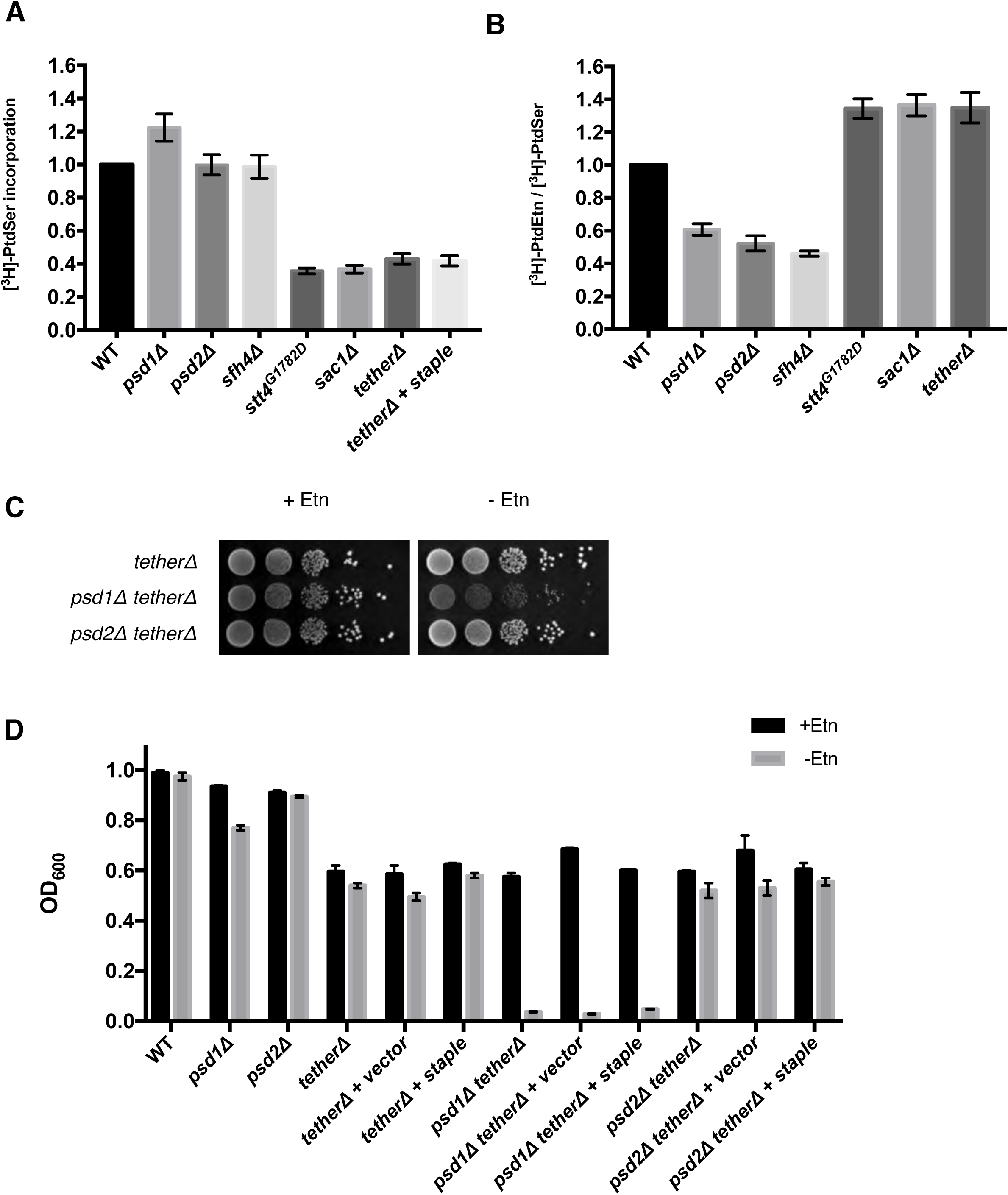
Complex relationship between PtdIns4*P* and PtdSer homeostasis and Psd2 activity in vivo. **(A)** Yeast strains of the indicated genotype were labeled with [^3^H]-serine for 3 hours and [^3^H]-PtdSer produced was resolved by TLC, scraped and quantified by scintillation counting. Total [^3^H]-PtdSer cpm for each strain was normalized by OD_600_ and related to the normalized [^3^H]-PtdSer level of WT cells (set at 1.0). Staple refers to expression of the artificial ER-plasma membrane tether. Values represent averages from at least three independent experiments and are plotted as mean ± SEM. **(B)** Yeast strains were labeled with [^3^H]-serine for 3 hours and the efficiency of PtdSer decarboxylation to PtdEtn was determined by TLC separation followed by scintillation counting. The conversion of PtdSer to PtdEtn was expressed as the [^3^H]-PtdEtn/[^3^H]-PtdSer ratios of the indicated strains and related to the [^3^H]-PtdEtn/[^3^H]-PtdSer ratio of the WT control analyzed in the same experiment. Values represent averages from at least three independent experiments and are plotted as mean ± SEM. **(C)** Isogenic yeast strains with the designated genotype were spotted in 10-fold dilution series on glucose minimal media with or without Etn supplementation (2mM) as indicated. Plates were incubated at 30°C for 48 hours. **(D)** Growth of the indicated yeast strains in glucose minimal SD liquid media with or without Etn (2mM). The assay was conducted as described in Figure 4B. Cultures were grown at 30°C for 46.5 hours and the final A_600_ of each culture was recorded. Staple refers to expression of the artificial ER-plasma membrane tether. Values represent averages of two independent experiments (n=2) plotted as mean ± range.

Whereas cells carrying individual *sfh4Δ, psd1Δ* or *psd2Δ* mutations showed undiminished incorporation of labeled amino acid into [^3^H]-PtdSer relative to WT, both the *stt4^G1782D^* and the *sac1Δ* mutants reproducibly exhibited significantly diminished incorporation (∼38% of WT; Figure 7A). Unexpectedly, those reductions in [^3^H]-PtdSer for *stt4^G1782D^*and *sac1Δ* mutants were not accompanied by decreased [^3^H]-PtdEtn/[^3^H]-PtdSer ratios -- indicating that neither *stt4^G1782D^*nor *sac1Δ* mutants reduced the efficiencies with which the PtdSer molecules that engaged the Psd2 pathway were converted to PtdEtn. This was in stark contrast to cells carrying individual *sfh4Δ, psd1Δ* or *psd2Δ* cells where [^3^H]-PtdEtn/[^3^H]-PtdSer ratios were decreased to ∼50% of WT values (Figure 7B). Those results indicated that: (i) contrary to a previous report that Psd2 is a minor contributor to PtdSer decarboxylation in yeast (Burgermeister, Birner-Grunberger et al., 2004), the Psd1 and Psd2 pathways contribute roughly equally to bulk cellular PtdSer decarboxylation capacity, and (ii) Sfh4 deficits compromise activity of the Psd2 pathway directly, whereas Stt4 and Sac1 insufficiencies do not.

### Stt4 and Sac1 deficits are recapitulated by functional ablation of ER-PM tethers

The metabolic phenotypes of *stt4^G1782D^* and *sac1Δ* mutants suggested derangements in how PtdSer synthesis is regulated or otherwise organized. The role of ER-plasma membrane (PM) contact sites as regulatory/organizing nodes for lipid synthesis has been noted (Quon, Sere et al., 2018). We therefore assessed ER-PM MCS status on the cellular PtdSer decarboxylation capacity. Thus, the *PSD1* or *PSD2* genes were deleted in a *tetherΔ* genetic background where the structural genes for six ER-PM membrane tethers (*TCB1, TCB2, TCB3, SCS2, SCS22, IST2*) are combinatorially deleted (Manford, Stefan et al., 2012). Indeed, *tetherΔ* cells recapitulated the phenotypes of *stt4^G1782D^* and *sac1Δ* mutants. That is, when interrogated for [^3^H]-serine incorporation into PtdSer, the *tetherΔ* mutant showed reduced efficiencies of incorporation that were only ∼40% of wild-type, *psd1Δ* or *psd2Δ* yeast (Figure 7A). In addition, whereas *sfh4Δ, psd1Δ* and *psd2Δ* mutants all exhibited reduced [^3^H]-PtdEtn/[^3^H]-PtdSer ratios relative to wild-type (Figure 7B), the *tetherΔ* mutant did not show any reduction at all – indicating that neither the Psd1 nor the Psd2 pathways were intrinsically compromised in the *tetherΔ* mutant.

Interestingly, *tetherΔpsd1Δ* cells displayed a partial Etn-auxotrophy characterized by no growth in Etn-free liquid media and weak growth on solid Etn-free media that was nonetheless superior to significantly tighter Etn auxotrophy of *stt4^G1782D^ psd1Δ* double mutants (Figure 7C,D, Suppl. Figure S7). This level of reduced ER-PM tether activity did not compromise the ability of the Psd1 pathway to fulfill cellular PtdSer decarboxylation requirements as *tetherΔpsd2Δ* cells were Etn prototrophs (Figure 7C,D). Thus, the sufficiency of the Psd2 pathway to meet cellular PtdSer decarboxylation requirements was selectively compromised in the *tetherΔ* mutant.

To determine whether simple tethering of ER to the PM was sufficient to rescue PtdSer homeostasis in *tetherΔ* cells, an artificial ER-PM protein tether was expressed in the double mutant (Quon, Sere et al., 2018). Expression of this artificial staple failed to rescue [^3^H]-PtdSer levels in *tetherΔ* yeast and failed to restore growth of *tetherΔpsd1Δ* cells in Etn-free liquid media (Figure 7D). That finding suggests these tethers, or a subset thereof, either play active roles in recruiting Sac1 or Stt4 to some functional membrane domain, or in regulating uptake of precursors for the Psd2 pathway. We interpret the selective compromise of Psd2 pathway capacity in the face of ER-PM tether deficiencies to reflect reduced PtdSer availability to Psd2 upon reductions in PtdSer synthesis.

### PtdSer decarboxylation is primarily required for de novo PtdCho biosynthesis

The compartmental segregation of the Psd1 and Psd2 pathways suggests these two pathways address unique cellular (compartment-specific?) PtdEtn requirements. Whereas we employed Etn auxotrophy in this study as phenotypic readout for Psd2 pathway activity, PtdSer metabolism also affects PtdCho biosynthesis (Boumann, Gubbens et al., 2006, Storey, Clay et al., 2001). The de novo pathway for PtdCho biosynthesis uses PtdEtn as precursor which, via a series of three headgroup methylation reactions, is converted to PtdCho. As the Etn-free media used throughout this study were also choline-deficient, contributions of PtdSer decarboxylation to de novo PtdCho synthesis could be assessed by addition of choline (rather than Etn) to the medium.

Indeed, supplementation of Etn-free growth medium with a low concentration of choline (10μM) was sufficient to partially rescue the growth defects of *psd1Δsfh4Δ* and of *psd1Δpsd2Δ* double mutants (Suppl. Figure S7). Notably, the Etn-prototrophy of these mutants in choline-supplemented media required a functional pathway for degradation of sphingosine-phosphate to Etn-phosphate catalyzed by the *DPL1* gene product. Even 100μM choline was insufficient to rescue growth of these mutants in a *dpl1Δ* genetic background (Suppl. Figure S7). Those data indicated that PtdSer decarboxylation in yeast primarily supports de novo PtdCho biosynthesis, and sharpen previous findings that the Etn-auxotrophy of *psd1Δpsd2Δ* double mutants is rescued in a Dpl1-dependent manner by excessive choline (Storey et al., 2001).

Strikingly, low choline (10μM) supplementation was more efficient in rescue of the growth defects of *psd1Δstt4^G1782D^, psd1Δsac1Δ* and *tetherΔ psd1Δ* cells relative to the rescue scored for *psd1Δsfh4Δ* and *psd1Δpsd2Δ* double mutants. The efficiencies of rescue were similar to those observed when growth medium was supplemented with 2mM Etn (Suppl. Figure S7). These data provide yet another indication that the contribution of Stt4-dependent PtdIns4*P* homeostasis to Psd2-mediated PtdEtn production is mechanistically distinct from that of Sfh4.

## DISCUSSION

The proposed ER-endosomal MCS that organizes Psd2-mediated decarboxylation offers an attractive experimental model for testing basic concepts that are presumed to apply generally to MCS biology. The uniqueness of this system lies in the fact that the proposed functional properties of this candidate MCS can be interpreted in terms of a clear physiological context (Voelker, 2005). The current Psd2 ER-endosomal MCS model rests on two lines of experimental evidence. First, genetic evidence identifies Sfh4 and Stt4 as important components as deficiencies in either function compromise Psd2 activity in vivo (Trotter et al., 1998, Wu, Routt et al., 2000). These data do not speak to whether the involvements of Sfh4 and/or Stt4 in the Psd2 pathway report direct or indirect effects. A second line of supporting biochemical evidence consists of an unvalidated chain of protein::protein interactions that connect Psd2 with Sfh4, the functionally uncharacterized Pbi1, and known MCS tethers of the VAP-family (Scs2 and Scs22) that bind Stt4 (Riekhof et al., 2014).

Herein, we show that the Sfh4 PITP and the Stt4 PtdIns 4-OH kinase are required for biologically-sufficient activation of Psd2 via a mechanism that requires Stt4 enzymatic activity and employs Sfh4 in what we describe as a non-canonical fashion. Indeed, the data identify Sfh4 as the only component other than Psd2 that can be confidently assigned as being directly involved in the yeast Psd2 pathway. The cumulative results report a substantially indirect role for Stt4 in regulating Psd2 activity. We presently favor a model where Stt4 mediates its indirect effects through PtdIns4*P* responsive control of bulk PtdSer synthesis and/or targeting of Psd2 to the correct intracellular compartment(s). Taken together, the data fail to support the foundational tenets of the Psd2 MCS model as proposed, and highlight the interpretive complexities associated with mechanistic dissection of MCS function.

### Re-evaluation of the existing Psd2 MCS model

Using genetic and metabolic tracer analyses, we find that neither the uncharacterized Pbi1, nor the MCS tethers Scs2 and Scs22, play any discernible role in regulating PtdSer decarboxylation by Psd2 in vivo. Similarly, neither functional ablation of each individual *OSH* gene, structural genes encoding other tethers (Vps13, Lam5, Lam6), nor individual overexpression of the purported PtdSer transfer proteins Osh6 and Osh7, had any significant biological consequences for Psd2 activity. Taken together, these collective data disqualify Scs2/Scs22, Pbi1, Vps13, Lam5, Lam6, and any individual member of the yeast *OSH* protein family as quantitatively significant regulators of Psd2 activity – at least under typical laboratory conditions. Sfh4 and Stt4 represent the only two individual components that exert significant control over Psd2-mediated PtdSer decarboxylation in cells.

### A Psd2::Sfh4::Stt4 metabolon?

When viewed in the context of the six yeast Sec14-like PITPs, the specific ability of Sfh4 to stimulate Psd2 activity can now be accounted for by the unique ability of Sfh4 to physically interact with Psd2 in vivo. The functional relevance of this interaction is convincingly demonstrated by the fact that the graded biochemical defects in the Sfh4:Psd2 interaction represented in an allelic series of Sfh4 variants were proportionately reflected in the severities of Psd2-dependent PtdSer decarboxylation defects exhibited by cells reconstituted for expression of those variants as sole sources of Sfh4.

Consistent with the genetic data identifying Stt4 as potentiator of Psd2 activity, a Psd2::Stt4 physical interaction was detected as well. This interaction was independent of Sfh4, and Sfh4 interaction with Psd2 was similarly refractory to diminished Stt4 activity. Given the lack of evidence for other components required for biologically sufficient Psd2 activity, we presently interpret the data to report direct Sfh4::Psd2 and Stt4::Psd2 interactions. While ultimate functional validation awaits the characterization of mutants that break the Psd2::Stt4 physical interaction, the collective data are consistent with the idea that Psd2, Sfh4 and Stt4 enter into a dedicated complex required for Psd2 activity in vivo. Such an arrangement supports proposals that specific PITPs engage PtdIns 4-OH kinases in dedicated signaling complexes (i.e. ‘pixels’) to effect specific biological outcomes (Grabon, Bankaitis et al., 2019). In this case, the data are consistent with assembly of what we describe as a Psd2 metabolon where the activities of a PITP and a PtdIns 4-OH kinase couple to the action of a ‘housekeeping’ metabolic enzyme.

### A non-canonical role for a PITP in regulating lipid metabolism

Given the established roles of PITPs in stimulating PtdIns 4-OH kinase activities in vivo (Bankaitis et al., 2010, Schaaf et al., 2008, Xie, Hur et al., 2018), the expected mechanism for how the Psd2::Sfh4::Stt4 metabolon operates was that the Sfh4 PITP employs its PtdIns-binding activities to potentiate Stt4 activity for production of a local PtdIns4*P* pool required for Psd2 activity. While our demonstration that *stt4^G1782D^* encodes a partially crippled enzyme which exhibits reduced PtdIns 4-OH kinase activity in vivo is consistent with this scenario, other data reject such a simple interpretation. Most notable in that regard are the surprising demonstrations that the PtdIns-binding/exchange activities of Sfh4 (and therefore its ability to stimulate PtdIns 4-OH kinase activity) were dispensable for Psd2 activation, and that Sfh4 activities in PtdIns-binding/exchange could be cleanly uncoupled from those required for promoting Psd2-dependent PtdSer decarboxylation.

While the residual low-level PtdIns-exchange activities of the Sfh4^T266D,K269A^ and Sfh4^T266W,K269A^ PtdIns-binding mutants might be sufficient to stimulate Stt4 activity in the close quarters of a Psd2-Sfh4-Stt4 metabolon, the available data do not support the case that Psd2 enzymatic activity is stimulated by PtdIns4*P* (see below). Rather, the data are most consistent with Sfh4 serving a scaffolding function required for Psd2 activation. Such a scaffolding function could serve to activate the decarboxylase in a non-catalytic subunit capacity. Alternatively, the Sfh4::Psd2 interaction might play a key role in ensuring the targeting of the decarboxylase to the appropriate intracellular compartment. In light of the observed Psd2::Stt4 physical interaction, the possibility that Stt4 might also contribute in a non-catalytic role to such a Psd2 localization strategy cannot be excluded. That specific scenario predicts breaking the Psd2::Stt4 interaction will elicit phenotypes that mimic those of Sfh4-deficient mutants.

### MCS tethers and homeostasis of Stt4-dependent PtdIns4*P* pools

Psd2 is a poorly expressed protein and Sfh4 and Stt4 are present in stoichiometric excess relative to Psd2 in vivo. While the Sfh4::Psd2 interaction is of clear physiological relevance, the genetic evidence for Stt4 involvement in Psd2 activity is difficult to interpret given the lack of in vivo data functionally validating the Psd2::Stt4 interaction. As Stt4 catalytic deficiencies apply to all Stt4 pools, not just to the Psd2-bound Stt4 pool, phenotypic data collected from *stt4^G1782D^* mutants sum both direct and indirect effects of Stt4 activity and PtdIns4*P* homeostasis on Psd2 metabolon activity.

The collective data indicate that, in addition to any direct functional role the Psd2::Stt4 interaction potentially serves, Stt4 also plays a significant and indirect role in modulating the Psd2 pathway in cells. The bulk of the supporting evidence to that effect comes from our paradoxical demonstration that attenuated Stt4-dependent PtdIns4*P* production (in *stt4^G1782D^* cells), large increases in Stt4-generated PtdIns4*P* pools due to compromised Sac1-phosphoinositide phosphatase-mediated degradation of (in *sac1Δ* cells), or combinatorial ablation of six known ER-plasma membrane tethers (*tetherΔ* cells), all evoked Etn (and Cho) auxotrophies in *psd1Δ* genetic backgrounds. Critically, these did so without affecting the efficiencies with which the PtdSer molecules that had entered the Psd2 pathway were decarboxylated. These disparate genetic backgrounds share two common features. First, bulk PtdSer synthesis was significantly reduced in *stt4^G1782D^, sac1Δ* and *tetherΔ* cells. Second, *stt4^G1782D^, sac1Δ* and *tetherΔ* mutants all exhibit significant derangements in PtdIns4*P* homeostasis. We show herein that production of this phosphoinositide is reduced in *stt4^G1782D^*yeast, while previous studies reported PtdIns4*P* accumulation to very high levels in *sac1Δ* and *tetherΔ* mutants (Guo et al., 1999, Manford et al., 2012, Rivas et al., 1999, Stefan et al., 2011).

The most parsimonious explanation of these results is that perturbed PtdIns4*P* homeostasis (either PtdIns4*P* deficit or excess) interferes with bulk PtdSer synthesis, and that the Psd2 pathway is particularly sensitive to reductions in the substrate pool. Several potential mechanisms account for such selective sensitivity of the Psd2 pathway to reduced PtdSer synthesis. First, if Psd1 has a lower apparent Km for PtdSer than does Psd2, the latter enzyme would be selectively disadvantaged under conditions where these decarboxylases compete for substrate from the same limiting pool. However, as Psd1 and Psd2 reside in distinct subcellular compartments, we consider it unlikely that the two enzymes compete directly for the same PtdSer pool. We find it a more reasonable conjecture that the compartment of Psd2 residence presents a membrane environment where accessible PtdSer becomes particularly limiting under conditions of reduced synthesis of bulk PtdSer. Given recent evidence that Psd1 is localized to both mitochondria and ER (i.e. the site of PtdSer synthesis) (Friedman, Kannan et al., 2018), it is plausible that Psd1 has more direct access to newly synthesized PtdSer than does Psd2.

Whereas reduced PtdSer synthesis alone could potentially account for attenuated Psd2 activity in *stt4^G1728D^*, *sac1Δ* and *tetherΔ* cells, the contributions of proper PtdIns4*P* homeostasis to biologically sufficient Psd2 activity are likely important as well. Given the recognized role of PtdIns4*P* as regulator of Golgi and endosomal membrane trafficking (Bankaitis et al., 2010, Graham & Burd, 2011, Wang, Mousley et al., 2019), a likely consequence of derangements in PtdIns4*P* homeostasis would be Psd2 mis-sorting to compartments whose membrane environments might be particularly hostile to biologically sufficient Psd2 activity under conditions of reduced PtdSer synthesis. Distinguishing between these possibilities demands that the intracellular localization of Psd2 be assigned at high-resolution, followed by subsequent determinations of whether localization of the enzyme is altered in the face of Sfh4 deficiencies or of deranged PtdIns4*P* homeostasis. Unfortunately, the vanishingly low cellular abundance of Psd2 frustrates attempts to assign a precise compartmental localization to the endogenous protein.

Finally, the data reported herein not only demand wholesale revision of the Psd2 MCS model, but also highlight the complexities associated with interpreting the types of data that are generally incorporated into mechanistic cartoons for MCS function. As one example, the non-canonical mechanism for PITP function described in this study emphasizes that caution be exercised in interpreting the functions of MCS-associated proteins that show lipid-transfer activities in vitro. MCS typically include at least one such component and it is almost universally assumed, in the absence of direct evidence, that such lipid-transfer activities translate to inter-organelle lipid-transfer in vivo. The indirect involvement of Stt4 and MCS tethers, in the face of genetic and/or interaction data that could be construed to argue otherwise, serves as another. Thus, the idea that MCS are involved in organizing lipid synthesis, rather than prosecuting inter-membrane lipid transport, deserves more serious consideration than is afforded to it in contemporary literature. In sum, this study emphasizes the necessity of direct assays, grounded in physiologically relevant contexts, for mechanistic dissections of MCS function.

## MATERIALS AND METHODS

### Yeast strains, media and reagents

Yeast media and standard genetic and molecular biology methods have been described (Cleves, McGee et al., 1991, Schaaf et al., 2008). All phospholipids were from Avanti Polar Lipids Inc. (Alabaster, AL), and restriction endonucleases from New England Biolabs (Ipswich, MA). All other standard reagents were obtained either from Sigma (St. Louis, MO) or Fisher Scientific (Norcross, GA). Yeast strain genotypes are listed in Supplemental Table 2 and plasmids are in Supplemental Table 3.

### [^3^H]-Serine metabolic radiolabeling and phospholipid analysis

Strains were grown in uracil-free minimal medium containing 3% glucose and 2 mM Etn to late-log phase at 30°C and shaking. Cultures were diluted to an OD_600_ = 0.3 and supplemented with 3.33 μCi/ml L-[^3^H] serine (ART 0246; American Radiolabeled Chemicals Inc., St. Louis, MO) and incubated for an additional 3h at 30°C and shaking. Incorporation of radiolabel was terminated by addition of ice-cold trichloroacetic acid (5% final concentration), and samples were incubated on ice for 30 min. Cell pellets were washed twice with cold ddH_2_O, resuspended in 1.5 ml ddH_2_O:absolute ethanol (1:4, v/v) and incubated at 100°C for 45 min. The lipids were then extracted from the 1:4 water:ethanol mixture by adding 4 ml of CHCl_3_, 4 ml of methanol and 3.3 ml of 0.2 M KCl, shaking, and centrifuging at top speed to separate the phases. The organic phase was collected and washed twice with 7.6 ml of the practical upper phase (PBS:methanol -- 9:10, v/v), saturated with CHCl3), dried under a stream of N_2_ gas and the lipid film was resuspended in CHCl_3_/CH_3_OH (2:1, v/v) charged with 1 mg/ml butylated hydroxytoluene. Lipids were resolved by Silica Gel H thin-layer chromatography (Analtech, Newark, DE) in a CHCl_3_/2-propanol/0.25% KCl/triethylamine (30:9:6:18, v/v/v/v) solvent system. After development, plates were sprayed with 0.2% (w/v) 8-anilino-1-naphthalenesulfonic acid and lipids were visualized under UV illumination. Individual lipid species were identified using internal standards (Avanti), and collected by scraping. Radioactivity was quantified by liquid scintillation counting.

### PtdIns4*P* quantification by thin-layer chromatography

Yeast strains were grown overnight in uracil-free minimal medium containing 3% glucose and 1% casamino acids (Cat #223050, Becton, Dickinson and Company, Franklin Lakes, NJ) and offered 10 μCi/ml [^3^H]*myo*-inositol (ART 0116A; American Radiolabeled Chemicals Inc., St. Louis, MO). After an incorporation window of at least 20h, labeling was terminated with trichloroacetic acid (5% final concentration) and samples were incubated on ice for 30 min. Cells were pelleted (16,000g for 1 min), washed twice in 500μl of cold ddH2O and resuspended in 500μl 4.5% perchloric acid. Approximately 300μl 0.5-mm glass beads were added, and cells were disrupted by vigorous agitation for 10 min in 1-min bursts with 1-min rest intervals on ice. Lysates from disrupted cells were collected and centrifuged at 16,000g for 10 min, the pellet washed with 500μl of 100mM EDTA (pH 7.4) and resuspended in 500μl of CHCl_3_/CH_3_OH/HCL (2:1:0.007). A two-phase separation was produced by addition of 100 μl of 0.6 M HCl, and samples vortexed for 5 min and centrifuged for 5 min (16,000g). The organic phase was collected, washed twice with 250μl of CH_3_OH/0.6 M HCl/CHCl_3_ (1:0.94:0.06), dried under N_2_ gas and resuspended in 50μl CHCl_3_3. Samples were resolved by thin-layer chromatography on Partisil LK6DF 60-Åsilica gel plates (Whatman, cat. no. 4866-821) using a CHCl_3_/CH_3_OH/ddH_2_O/NH_4_OH (1:0.83:0.15:0.1) solvent system. Radiolabeled lipids were visualized by autoradiography.

### Fluorescence imaging

Yeast cultures for GFP-2xPH^Osh2^ imaging were grown in synthetic defined medium lacking uracil at 30°C with shaking. Cells were collected from liquid cultures grown in uracil-free YNB supplemented containing 3% glucose and 1% casamino acids at 25°C by centrifugation at 5,000g for 1 min and resuspended into fresh uracil-deficient medium before analysis. Cells were sealed under a coverslip and examined at 25°C. The imaging system employed a CFI plan Apochromat Lambda 100x NA 1.45 oil immersion objective lens mounted on a Nikon Ti-U microscope base (Nikon, Melville, NY) interfaced to a Photometrics CoolSNAP HQ2 high-sensitivity monochrome CCD camera (Roper Scientific, Ottobrunn, Germany) or an Andor Neo sCMOS CCD camera (Andor Technology, Belfast, UK). A Lumen 200 Illumination System (Prior Scientific Inc., Rockland, MA.) was used in conjunction with a B-2E/C (465–495 nm/515–555 nm; EX/EM) filter set (Nikon, Melville, NY). Images were captured using the Nikon NIS Elements software package (Nikon, Melville, NY, version 4.10). Figures were generated using Adobe Photoshop 7.0.1.

### Homology modeling

Structural models of Sfh4 were generated with MOE 2013.08 Modeling package. The target Sfh4 sequence was aligned to the PtdIns-bound Sfh3 structure (PDB I.D. 4J7Q; 1.55 Å resolution) with a sequence similarity of ∼66%. PtdIns ligand was included in the environment to generate induced fit homology models of Sfh4-PtdIns complex. By default, 10 independent intermediate models were generated. These different intermediate homology models were generated as a result of permutational selection of different loop candidates and side chain rotamers. The intermediate model, which scored best according to the selected force field (Amber99), was chosen as the final model and was subjected to further optimization.

### Protein purification

pET28b plasmid derivatives programming expression of the appropriate recombinant proteins were transformed into BL21 *E. coli* (DE3; New England BioLabs Inc, Ipswich, MA). Recombinant proteins of interest were bound to TALON metal affinity beads (Clontech, Mountain View, CA) and eluted with imidazole (gradient 10–200 mM) and dialyzed (prod. no. 68100, Thermo Scientific, Rockford, IL). In the case of Sec14, dialysis was against 300 mM NaCl, 25 mM Na_2_HPO_4_ (pH = 7.5), 5 mM β-mercaptoethanol. Purified Sfh4 proteins were dialyzed against the same buffer with the exception that 50 mM Na_2_HPO_4_ was used. Protein mass was quantified by SDS-PAGE using BSA standards and A_280_ measurements.

### PtdIns-transfer assays

Purified recombinant proteins were preincubated in the presence of [^3^H] *myo*-inositol labeled acceptor membranes, in buffer (300 mM NaCl, 25mM Na_2_HPO4, pH 7.5) for 30 min at 37°C before initiating the assay by addition of radiolabeled donor membranes (Bankaitis, Aitken et al., 1990, Schaaf et al., 2008).

### Real-time fluorescence dequenching PyrPtdIns-transfer assays

Measurements of protein-mediated PyrPtdIns-transfer activity from donor vesicles (with TNP-PtdEtn quencher) to acceptor vesicles (without TNP-PtdEtn quencher) were conducted using a real-time fluorescence dequenching assay as described previously (Grabon, Orlowski et al., 2017, Huang, Ghosh et al., 2016). Fluorescence intensity (excitation 346 nm; emission 378 nm) was recorded as a function of time at 37°C using a Horiba Fluorolog 3 fluorimeter (Horiba, Kyoto, Japan). To titrate PtdIns-transfer activities, 1μg aliquots of purified recombinant protein were serially injected into assay mixtures at 250s time intervals (5 injections total).

### Co-Immunoprecipitation experiments

All assays were performed using lysed spheroplasts as described previously (Gulshan et al., 2010). Cultures (100 ml) were grown to early log phase at 30°C and shaking. Cells were collected by centrifugation and washed in spheroplast solution I (1M sorbitol, 10mM MgCl_2_, 50mM K_2_HPO_4_, 30mM dithiothreitol (DTT), 1mM PMSF). Cells were then resuspended in spheroplast solution II (1M sorbitol, 10mM MgCl_2_, 50mM K_2_HPO_4_, 30mM DTT, 1mM PMSF, 25mM sodium succinate, 1mM PMSF, 250 U/ml lyticase) and incubated at 37°C for 15-30 min on a shaker at 100 rpm.

Spheroplasts were collected by centrifugation and disrupted in lysis buffer (1% Triton X-100, 0.15M NaCl, 50mM Tris-HCl, 2mM EDTA, 200uM sodium vanadate, 1mM PMSF and protease inhibitor tablets) with vigorous vortexing with 0.5 mm glass beads. Cell-free extracts were clarified by centrifuging lysates at 20,000 xg for 5 min. Supernatants were collected and lysate protein concentrations determined by BCA assay for purposes of normalization. For immunoprecipitation, cell lysates (∼12 mg total protein) were incubated with anti-TAP antibody (ThermoFisher #CAB1001) for 4 h at 4°C. Complexes were captured by addition of Protein A/G-magnetic beads (ThermoFisher, Pierce Protein A/G Magnetic beads, CAT#88802) reconstituted in lysis buffer to the samples followed by incubation for 2 h at 4°C. Protein A/G-magnetic beads were subsequently recovered on a magnetic field stand and washed 4 times in wash buffer (0.15M NaCl, 50mM Tris-HCl, 2mM EDTA, 200uM sodium vanadate, 1mM PMSF and protease inhibitor tablets). Bound complexes were eluted from the beads by adding Laemmli buffer and heating. Precipitated materials were resolved by SDS-PAGE and immunoblotted with anti-HA antibody (BioLegend Cat# 901501) and anti-TAP antibody (ThermoFisher CAB1001). Immunoblot images were acquired using Bio-Rad ChemiDoc™ XRS+ system.

## Supporting information

Combined Supplementary Materials

## ACKNOWLEDGEMENTS

We are particularly grateful to W. Scott Moye-Rowley (Department of Molecular Medicine, Univ. Iowa) for his generous gifts of yeast strains and plasmids, and for his helpful comments on the manuscript. This work was supported by grants from the National Institutes of Health (R01 GM44530; R35 GM131804) to VAB. The MS laboratory is supported by an SFB1190 (Compartmental gates and contact sites) of the DFG and a VolksWagen Life grant. MS is an incumbent of the Dr. Gilbert Omenn and Martha Darling Professorial Chair in Molecular Genetics. MEB is supported by a PhD fellowship from the Azrieli Foundation. The authors declare no financial conflicts.

