## Supplementary material for "Non-Canonical Regulation of Phosphatidylserine Metabolism by a Phosphatidylinositol Transfer Protein and a Phosphatidylinositol 4-OH Kinase": Combined Supplementary Materials

#### Legends to Supplementary Figures

**Figure S1. Design of the synthetic gene array screen for identifying components required for Psd2 activity in vivo.** Schematic illustration of the screen aimed to find non-essential genes whose functional ablation, when combined with the *psd1Δ* allele, retards growth on Etn-free media. A *psd1Δ* allele was recombined into the yeast gene deletion collection using a standard automated mating, sporulation and selection protocol. Colony sizes of yeast containing both the query library gene deletion and *psd1Δ* (double mutants, DM) were analyzed by relating colony sizes achieved on plates lacking Etn relative to those on Etn-replete medium. To validate that the observed growth deficit was Δ*psd1*-dependent, and not simply a result of the library background, the same analysis was performed on yeast that were only selected for the mutant allele from the library (single mutants, SM). Normalized growth deficits were calculated and DM strains that exhibited a growth score < 1 standard deviation from the mean (<0.47) were considered as potential hits.

**Figure S2. Phenotypic assays of *stt4* mutants and Stt4 subunit over-expressing cells.** **(A)** Isogenic $stt4^{ts}$ yeast transformed with YCp(*URA3*) or derivatives driving individual expression of *STT4*, $stt4^{G1782D}$ or $stt4^{D1752A}$ from the natural *STT4* promoter were spotted in 10-fold dilution series on uracil-free media and incubated for 48 hours at the permissive- and restrictive temperatures of 30$^{o}$C and 37$^{o}$C. **(B)** Isogenic *psd1Δ* $stt4^{G1782D}$ yeast transformed with YCp(*URA3*) or derivatives driving individual expression of *STT4*, $stt4^{G1782D}$ or $stt4^{D1752A}$ genes from the natural *STT4* promoter were spotted in 10-fold dilution series on uracil-free media +/- 2 mM Etn and incubated at 30°C for 48 hours. **(C)** Isogenic yeast strains of the designated genotype were spotted in 10-fold dilution series on synthetic complete (SC) media +/- 2 mM Etn and incubated at 30°C for 48 hours. **(D)** Isogenic *psd1Δ* or *psd1Δ stt4*$^{G1782D}$ yeast transformed with multicopy YEp vectors driving *YPP1*, *EFR3* or *SFK1* over-expression (from their respective natural promoters) were spotted in 10-fold dilution series on uracil-free media +/- 2 mM Etn and incubated at 30°C for 48 hours.

**Figure S3.** **Sfh4::PtdIns homology model and the PtdIns-binding motif.** **(A)** Isogenic *psd1Δ sfh4Δ* yeast transformed with yeast episomal *URA3* plasmids driving high levels of expression of each individual member of the yeast Sec14-like PITP family (identified at left), from the *PMA1* promoter, were spotted in a 10-fold dilution series on uracil-free selection media with or without Etn as indicated at top. Plates were incubated at 30°C for 48 hours. (**B**) Figure illustrates homology model of Sfh4::PtdIns complex (gray ribbon) threaded onto the Sfh3 crystal structure (PDB i.d 4J7Q) template (orange ribbon). **(C)** Sfh4::PtdIns homology model (gray ribbon) is superimposed onto the Sfh1 structure (PDB i.d 3B7N, brown ribbon). Figure shows conservation of structural fold and of the PtdIns binding pose. **(D)** Homology model of the Sfh4::PtdIns complex is rendered as gray ribbon. Enlarged panel highlights the conserved Sec14-like PITP PtdIns headgroup coordination motif, and illustrate conserved residues $T_{266}$ and $K_{269}$ interacting with PtdIns head group phosphate via a network of hydrogen bonds. **(E)** Fluorescence dequenching PtdIns-transfer assay for purified recombinant Sfh4 and Sfh4$^{266D,K269A}$. Fluorescence intensity of pyrene-PtdIns is plotted as a function of time. Black arrows identify points at which a 1 ug increment of the indicated protein was added. After protein addition, the observed increase in fluorescence intensity is directly proportional to the relative transfer efficiency of the protein being assayed. Values represent averages from two independent experiments plotted as mean ± standard deviation.

**Figure S4.** **PtdIn4*P* phosphatase and Sfh4/Stt4 requirement for Psd2 function. (A)** Yeast growth assay showing that *sac1Δ* gene failed to restore Psd2 pathway activity in mutants ablated for Sfh4 function and compromised for Stt4 activity. Isogenic *psd1Δ, psd1Δ* $stt4^{G1782D}$, *psd1Δ* $stt4^{G1782D}$ *sac1Δ, psd1Δ sfh4Δ, psd1Δ sfh4Δ sac1Δ* yeast were spotted in 10-fold dilution series on SC media +/- 2 mM Etn and incubated at 30°C for 48 hours. **(B)** Yeast growth assay showing transplacement of *inp51Δ, inp52Δ, inp53Δ, inp54Δ* alleles into a *psd1Δ sfh4Δ* strain failed to rescue Psd2 activity. Isogenic yeast strains with the designated genotype were spotted in 10-fold dilution series on SC media +/- 2 mM Etn and incubated at 30°C for 48 hours.

**Figure S5.** **The mutagenic PCR screen and the fluorescence dequenching PtdIns-transfer assay of *sfh4* mutants. (A)** Design of the mutagenic PCR screen for isolation of Sfh4 mutants specifically defective in Psd2 activity. A PCR-mutagenized *SFH4* fragment library was co-transformed with a gapped YCp(*URA3*) plasmid into the *ura3 sfh4Δ psd1Δ* $sec14^{ts}$ recipient strain. Transformants displaying the unselected parental Etn auxotrophy and the ability to grow at 37°C were identified. **(B)** Fluorescence dequenching PtdIns-transfer assay for purified recombinant Sfh4, $Sfh4^{F175L}$, $Sfh4^{F175A}$, $Sfh4^{F175R}$ and $Sfh4^{F175E}$ proteins. Fluorescence intensity of pyrene-PtdIns is plotted as a function of time. Black arrows identify points at which a 1 ug increment of the indicated protein was added. After protein addition, the observed increase in fluorescence intensity is directly proportional to the relative transfer efficiency of the protein being assayed. Values represent averages from two independent experiments plotted as mean ± standard deviation. These mutant Sfh4 data were obtained in the same experiments as the data reported in Suppl. Figure S3E, so the same Sfh4 control data are represented in both Figures.

**Figure S6. Co-Precipitation of Stt4 with Psd2. (A)** Wild-type yeast cells, where *STT4-TAP* or *SFH3-TAP* cassettes were transplaced into the corresponding gene loci, were transformed with a multicopy YEp plasmid driving Psd2-HA expression. Transformants were cultured to mid-log phase and cell-free lysates prepared. Co-precipitation experiments were performed as described in the legend to Figure 5A. **(B)** Wild-type or *psd2Δ* yeast cells with *STT4-TAP*, *stt4*$^{G1782D}$*-TAP* or *PIK1-TAP* cassettes transplaced for the endogenous genes were transformed with a YEp yeast episomal plasmid expressing Psd2-HA, were cultured to mid-log phase and cell-free lysates prepared. Co-IP experiment was performed as described in the legend to Figure 5A.

**Figure S7. Relationship between PtdSer decarboxylation pathways and de novo PtdCho biosynthesis.** Isogenic yeast strains with the designated genotype were spotted in 10-fold dilution series on SC media +2 mM Etn, +10 uM choline, +100 uM choline or without Etn or choline and incubated at 30°C for 48 hours.

### Figure S1

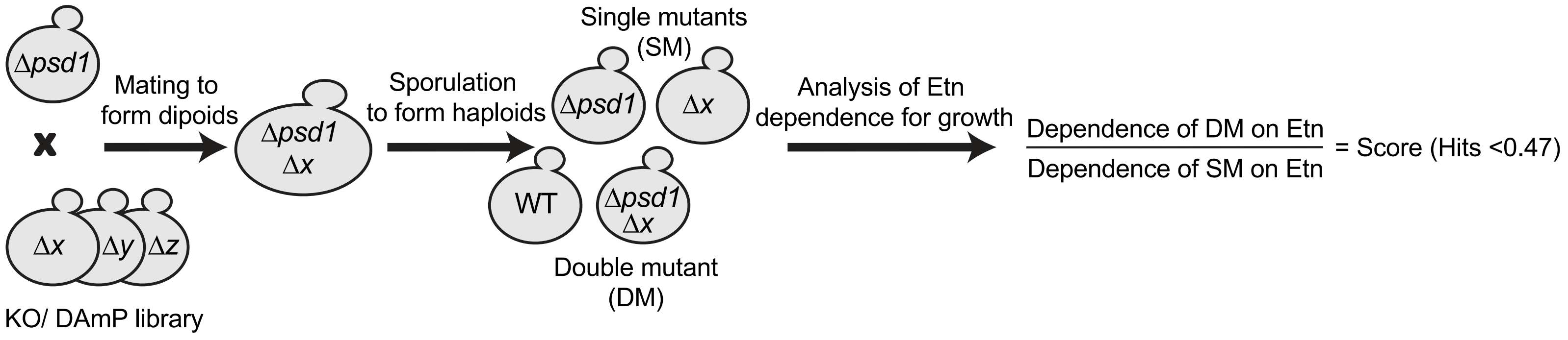
Δpsd1
x
Δx
Δy
Δz
KO/ DAmP library
Mating to form dipoids
Δpsd1
Δx
Sporulation to form haploids
Single mutants (SM)
Δpsd1
Δx
WT
Δpsd1
Δx
Double mutant (DM)
Analysis of Etn dependence for growth
Dependence of DM on Etn
Dependence of SM on Etn
= Score (Hits <0.47)

### Figure S2

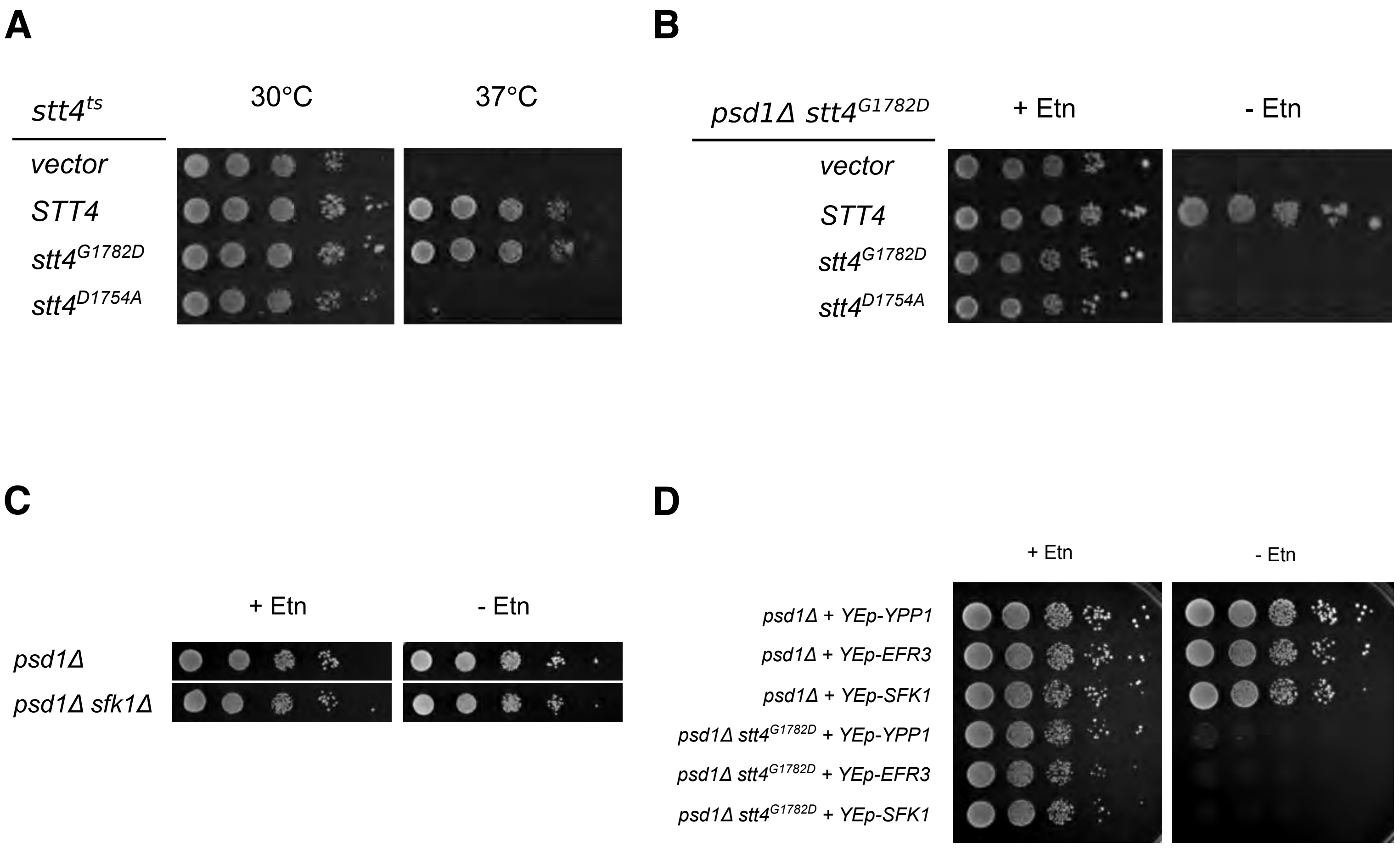
A
stt4ts
30°C
37°C
vector
STT4
stt4G1782D
stt4D1754A
B
psd1Δ stt4G1782D
+ Etn
- Etn
vector
STT4
stt4G1782D
stt4D1754A
C
+ Etn
- Etn
psd1Δ
psd1Δ sfk1Δ
D
+ Etn
- Etn
psd1Δ + YEp-YPP1
psd1Δ + YEp-EFR3
psd1Δ + YEp-SFK1
psd1Δ stt4G1782D + YEp-YPP1
psd1Δ stt4G1782D + YEp-EFR3
psd1Δ stt4G1782D + YEp-SFK1

### Figure S3

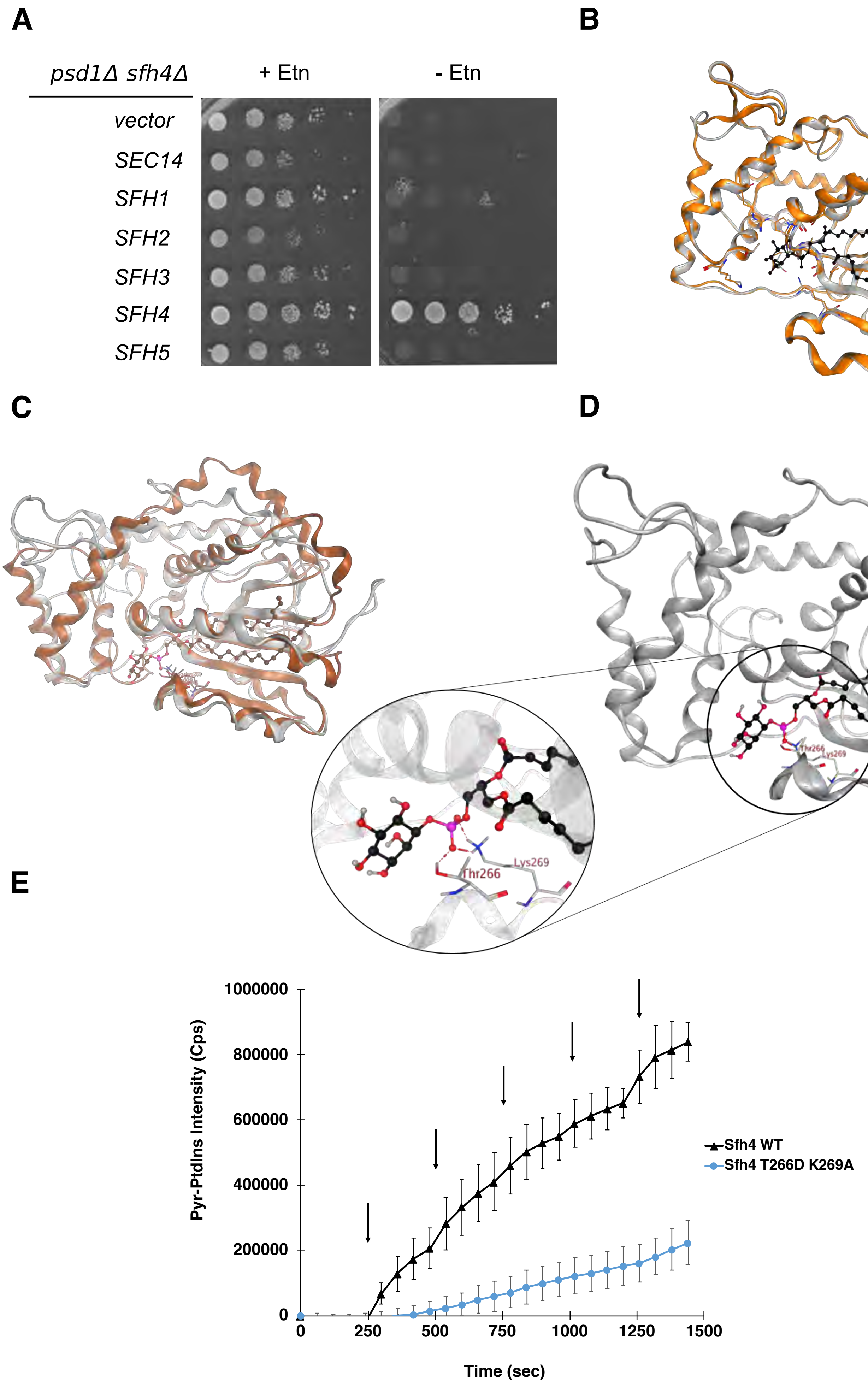
A
B
C
D
E
psd1Δ sfh4Δ
+ Etn
- Etn
vector
SEC14
SFH1
SFH2
SFH3
SFH4
SFH5
Thr266
Lys269
Pyr-PtdIns Intensity (Cps)
1000000
800000
600000
400000
200000
0
0
250
500
750
1000
1250
1500
Time (sec)
Sfh4 WT
Sfh4 T266D K269A

### Figure S4

**A**

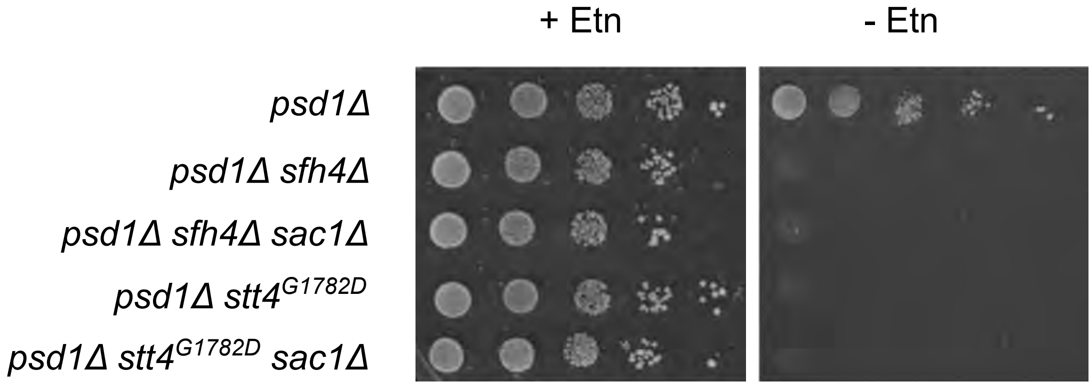
+ Etn
- Etn
psd1Δ
psd1Δ sfh4Δ
psd1Δ sfh4Δ sac1Δ
psd1Δ stt4$^{G1782D}$
psd1Δ stt4$^{G1782D}$ sac1Δ

**B**

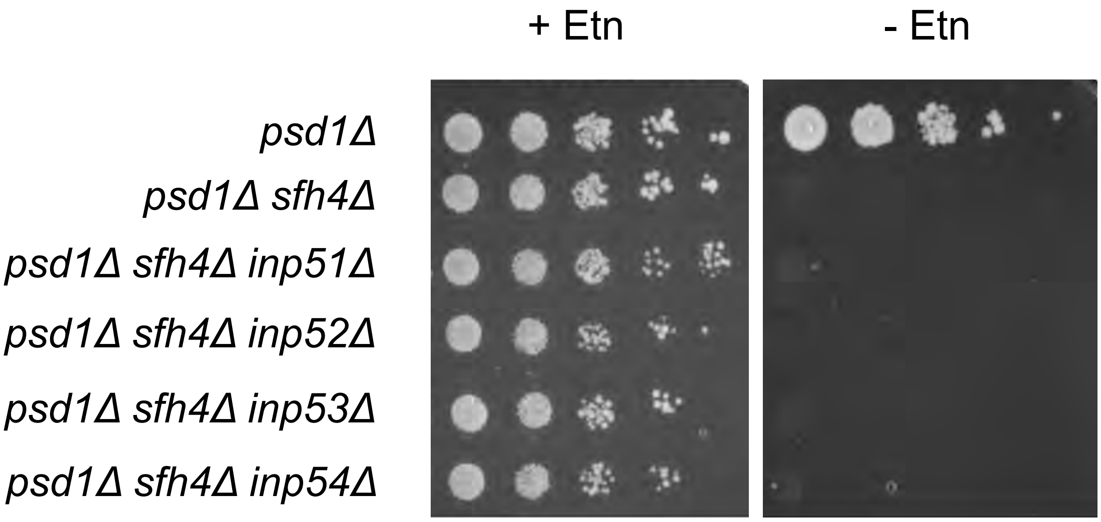
+ Etn
- Etn
psd1Δ
psd1Δ sfh4Δ
psd1Δ sfh4Δ inp51Δ
psd1Δ sfh4Δ inp52Δ
psd1Δ sfh4Δ inp53Δ
psd1Δ sfh4Δ inp54Δ

### Figure S5

**A**

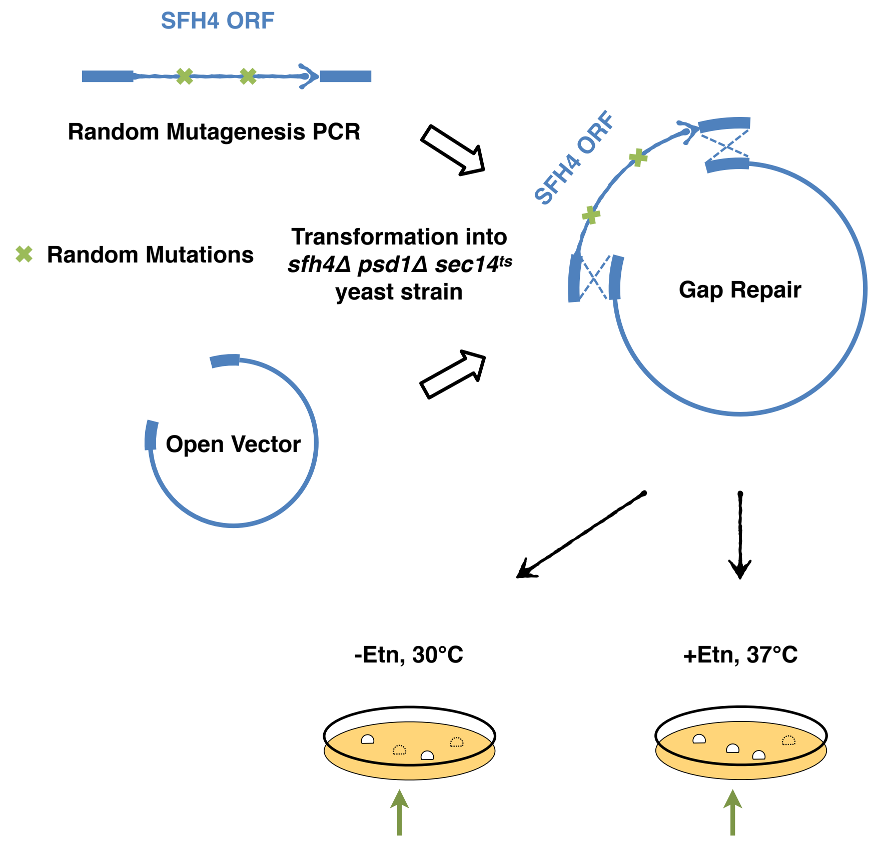
SFH4 ORF
Random Mutagenesis PCR
Random Mutations
Transformation into
*sfh4Δ psd1Δ sec14ts*
yeast strain
SFH4 ORF
Gap Repair
Open Vector
-Etn, 30°C
+Etn, 37°C

**B**

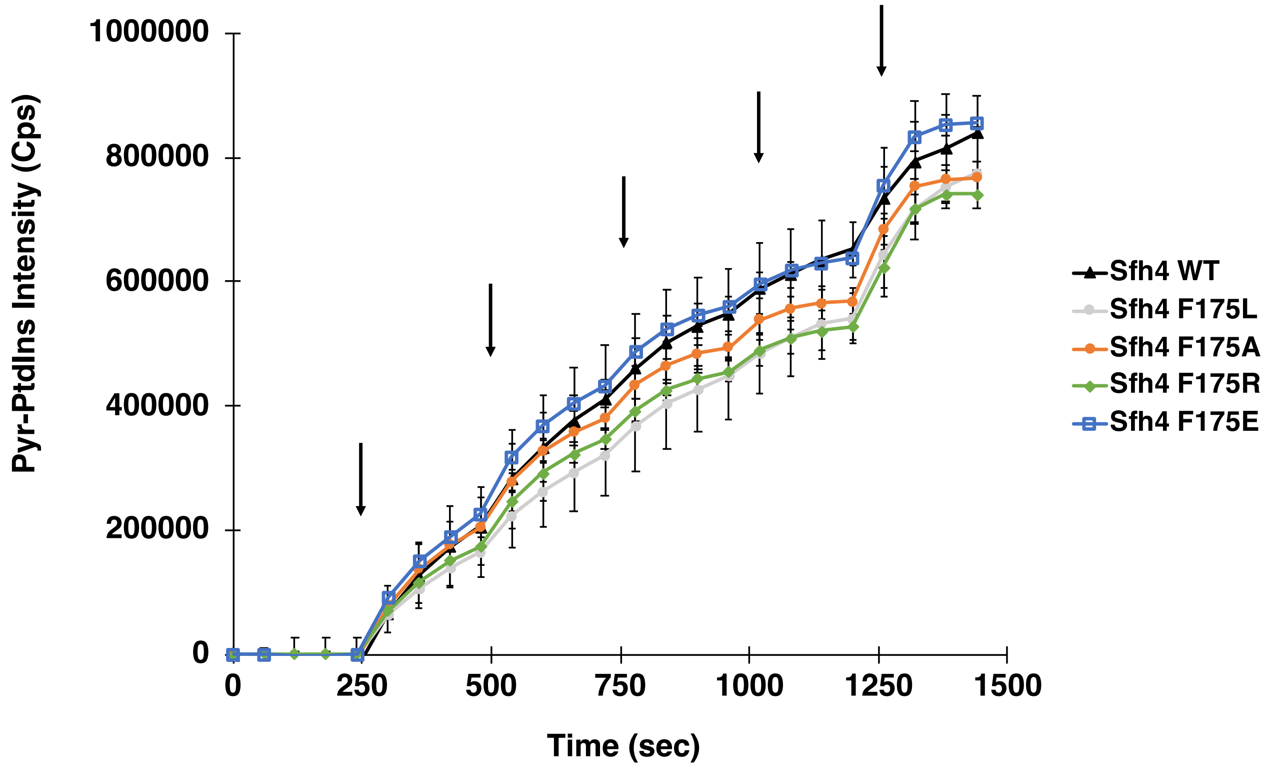
1000000
800000
600000
400000
200000
0
Pyr-PtdIns Intensity (Cps)
0
250
500
750
1000
1250
1500
Time (sec)
Sfh4 WT
Sfh4 F175L
Sfh4 F175A
Sfh4 F175R
Sfh4 F175E

### Figure S6

**A**

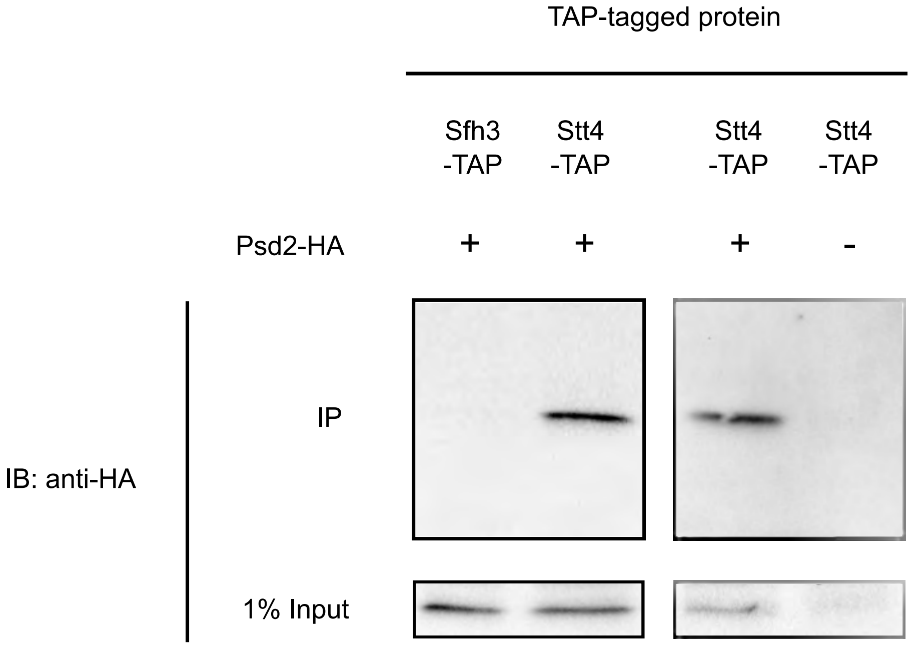
TAP-tagged protein
Sfh3 -TAP
Stt4 -TAP
Stt4 -TAP
Stt4 -TAP
Psd2-HA
+
+
+
-
IB: anti-HA
IP
1% Input

**B**

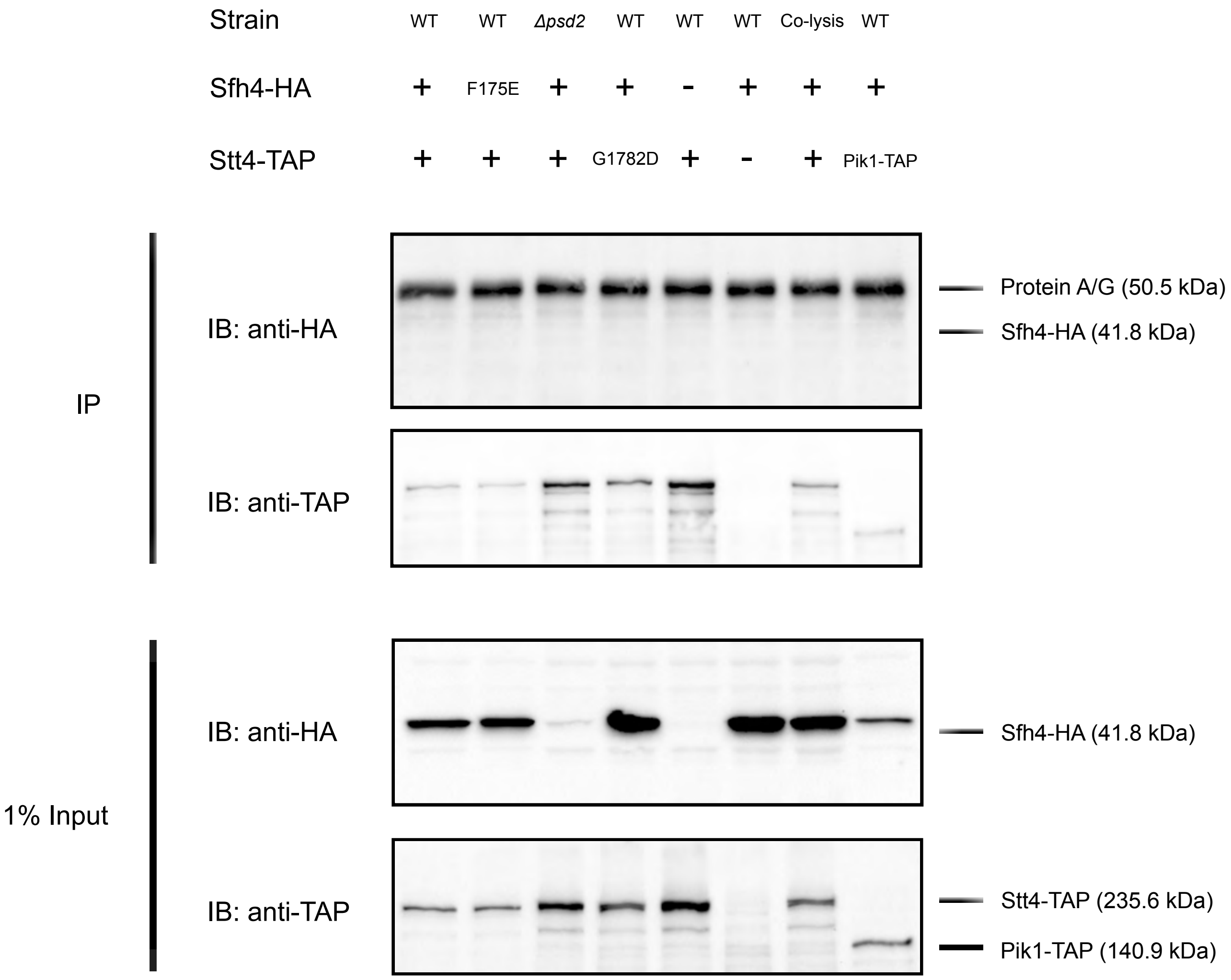
Strain
WT
WT
Δpsd2
WT
WT
WT
Co-lysis
WT
Sfh4-HA
+
F175E
+
+
-
+
+
+
Stt4-TAP
+
+
+
G1782D
+
-
+
Pik1-TAP
IP
IB: anti-HA
Protein A/G (50.5 kDa)
Sfh4-HA (41.8 kDa)
IB: anti-TAP
1% Input
IB: anti-HA
Sfh4-HA (41.8 kDa)
IB: anti-TAP
Stt4-TAP (235.6 kDa)
Pik1-TAP (140.9 kDa)

### Figure S7

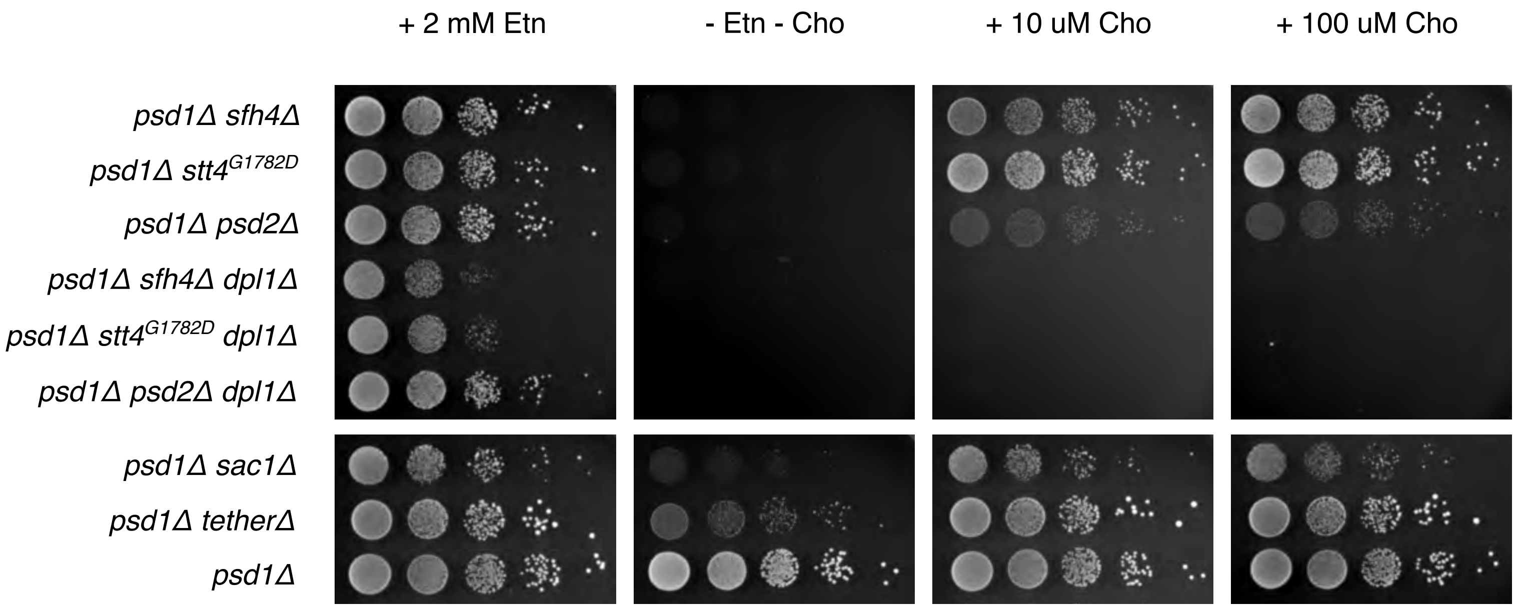

+ 2 mM Etn
- Etn - Cho
+ 10 uM Cho
+ 100 uM Cho
*psd1Δ sfh4Δ*
*psd1Δ stt4$^{G1782D}$*
*psd1Δ psd2Δ*
*psd1Δ sfh4Δ dpl1Δ*
*psd1Δ stt4$^{G1782D}$ dpl1Δ*
*psd1Δ psd2Δ dpl1Δ*
*psd1Δ sac1Δ*
*psd1Δ tetherΔ*
*psd1Δ*

#### Legends to Tables

**Table S1. A Synthetic Genetic Arrays (SGA) screen interrogating yeast genome for the loss-of-function (LOF) of any of yeast non-essential gene(s) which deactivates Psd2 pathway.** The list of hits from the screen performed to identify non-essential genes whose deletion, in combination with a *psd1Δ* allele, resulted in reduced colony size when cultured on Etn-free media (see Suppl. Figure S1). SM= Single mutants, DM= Double mutants. Strains with colony size smaller than 50, when grown on Etn-replete medium, were considered as background and were omitted.

**Supplementary Table S1. Genes whose deletion/hypomorphic allele in combination with a *psd1 Δ* allele resulted in a growth delay on media without ethanolamine (Etn)**

| ORF | Gene | Description | Area SM+Etn | Area SM-Etn | Area DM+Etn | Area DM-Etn | DM-Etn/DM+Etn | SM-Etn/SM+Etn | (DM-Etn/DM+Etn)/(SM-Etn/SM+Etn) |
|---|---|---|---|---|---|---|---|---|---|
| YNL264C | SFH4 | Phosphatidylinositol transfer protein (PITP); downregulates Plb1p-media | 185 | 125 | 75 | 0 | 0.00 | 0.68 | 0.00 |
| YGR170W | PSD2 | Phosphatidylserine decarboxylase of the Golgi and vacuolar membranes | 160 | 120 | 146 | 18 | 0.12 | 0.75 | 0.16 |
| YDL006W | PTC1 | Type 2C protein phosphatase (PP2C); dephosphorylates Hog1p, inactivat | 160 | 325 | 208 | 118 | 0.57 | 2.03 | 0.28 |
| YDR322C- | TIM11 | Subunit e of mitochondrial F1F0-ATPase; ATPase is a large, evolutionaril | 164 | 181 | 81 | 25 | 0.31 | 1.10 | 0.28 |
| YIL123W | SIM1 | Protein of the SUN family (Sim1p, Uth1p, Nca3p, Sun4p); may participat | 171 | 150 | 371 | 93 | 0.25 | 0.88 | 0.29 |
| YIL162W | SUC2 | Invertase; sucrose hydrolyzing enzyme; a secreted, glycosylated form is | 189 | 386 | 210 | 125 | 0.60 | 2.04 | 0.29 |
| YNL124W | NAF1 | RNA-binding protein required for the assembly of box H/ACA snoRNPs; t | 211 | 141 | 277 | 55 | 0.20 | 0.67 | 0.30 |
| YLR087C | CSF1 | Protein required for fermentation at low temperature; the authentic, no | 94 | 86 | 97 | 27 | 0.28 | 0.91 | 0.30 |
| YIL121W | QDR2 | Plasma membrane transporter of the major facilitator superfamily; mem | 164 | 127 | 365 | 88 | 0.24 | 0.77 | 0.31 |
| YNL050C | YNL050C | Putative protein of unknown function; YNL050c is not an essential gene | 215 | 473 | 167 | 117 | 0.70 | 2.20 | 0.32 |
| YBR137W | YBR137W | Protein of unknown function; localized to the cytoplasm; binds to Replic | 197 | 152 | 387 | 97 | 0.25 | 0.77 | 0.32 |
| YDR394W | RPT3 | ATPase of the 19S regulatory particle of the 26S proteasome; one of ATP | 194 | 135 | 301 | 69 | 0.23 | 0.70 | 0.33 |
| YDL095W | PMT1 | Protein O-mannosyltransferase of the ER membrane; transfers mannose | 163 | 135 | 310 | 90 | 0.29 | 0.83 | 0.35 |
| YLR262C-/ | TMA7 | Protein of unknown that associates with ribosomes; null mutant exhibits | 139 | 106 | 62 | 17 | 0.27 | 0.76 | 0.36 |
| YIL040W | APQ12 | Protein required for nuclear envelope morphology; nuclear pore comple | 79 | 111 | 75 | 38 | 0.51 | 1.41 | 0.36 |
| YDR438W | THI74 | Mitochondrial transporter repressible by thiamine; THI74 has a paralog, | 170 | 138 | 353 | 104 | 0.29 | 0.81 | 0.36 |
| YBR058C-. | TSC3 | Protein that stimulates the activity of serine palmitoyltransferase; invol | 139 | 148 | 87 | 34 | 0.39 | 1.06 | 0.37 |
| YNL059C | ARP5 | Nuclear actin-related protein involved in chromatin remodeling; compor | 82 | 97 | 69 | 30 | 0.43 | 1.18 | 0.37 |
| YFL031W | HAC1 | Basic leucine zipper (bZIP) transcription factor (ATF/CREB1 homolog); re | 146 | 164 | 115 | 49 | 0.43 | 1.12 | 0.38 |
| YBR138C | YBR138C | Cytoplasmic protein of unknown function; potentially phosphorylated by | 155 | 184 | 181 | 82 | 0.45 | 1.19 | 0.38 |
| YDR202C | RAV2 | Subunit of RAVE complex (Rav1p, Rav2p, Skp1p); the RAVE complex assc | 161 | 130 | 254 | 79 | 0.31 | 0.81 | 0.39 |
| YDR080W | VPS41 | Vacuolar membrane protein that is a subunit of the HOPS complex; esse | 108 | 64 | 83 | 19 | 0.23 | 0.59 | 0.39 |
| YNR012W | URK1 | Uridine/cytidine kinase; component of the pyrimidine ribonucleotide sal | 161 | 346 | 143 | 120 | 0.84 | 2.15 | 0.39 |
| YKL028W | TFA1 | TFIIE large subunit; involved in recruitment of RNA polymerase II to the | 173 | 139 | 282 | 89 | 0.32 | 0.80 | 0.39 |
| YBR029C | CDS1 | Phosphatidate cytidylyltransferase (CDP-diglyceride synthetase); an enz | 179 | 155 | 295 | 102 | 0.35 | 0.87 | 0.40 |
| YFL031W | HAC1 | Basic leucine zipper (bZIP) transcription factor (ATF/CREB1 homolog); re | 144 | 160 | 81 | 36 | 0.44 | 1.11 | 0.40 |
| YHR079C | IRE1 | Serine-threonine kinase and endoribonuclease; transmembrane protein | 132 | 116 | 125 | 44 | 0.35 | 0.88 | 0.40 |
| YKL212W | SAC1 | Phosphatidylinositol phosphate (PtdInsP) phosphatase; involved in hydr | 114 | 99 | 66 | 23 | 0.35 | 0.87 | 0.40 |
| YLR276C | DBP9 | DEAD-box protein required for 27S rRNA processing; exhibits DNA, RNA | 213 | 133 | 327 | 83 | 0.25 | 0.62 | 0.41 |
| YNL041C | COG6 | Component of the conserved oligomeric Golgi complex; a cytosolic tethe | 161 | 106 | 183 | 49 | 0.27 | 0.66 | 0.41 |
| YDR444W | YDR444W | Putative hydrolase acting on ester bonds | 175 | 242 | 142 | 81 | 0.57 | 1.38 | 0.41 |
| YGL140C | YGL140C | Putative protein of unknown function; non-essential gene; contains mul | 203 | 220 | 205 | 92 | 0.45 | 1.08 | 0.41 |
| YKR031C | SPO14 | Phospholipase D; catalyzes the hydrolysis of phosphatidylcholine, produ | 118 | 182 | 97 | 62 | 0.64 | 1.54 | 0.41 |
| YGR204W | ADE3 | Cytoplasmic trifunctional enzyme C1-tetrahydrofolate synthase; involve | 179 | 184 | 248 | 106 | 0.43 | 1.03 | 0.42 |
| YPR079W | MRL1 | Membrane protein; has similarity to mammalian mannose-6-phosphate | 201 | 314 | 175 | 114 | 0.65 | 1.56 | 0.42 |
| YML049C | RSE1 | Protein involved in pre-mRNA splicing; component of the pre-spliceoson | 203 | 122 | 330 | 83 | 0.25 | 0.60 | 0.42 |
| YML085C | TUB1 | Alpha-tubulin; associates with beta-tubulin (Tub2p) to form tubulin dime | 199 | 139 | 372 | 109 | 0.29 | 0.70 | 0.42 |
| YEL036C | ANP1 | Subunit of the alpha-1,6 mannosyltransferase complex; type II membrar | 202 | 184 | 278 | 107 | 0.38 | 0.91 | 0.42 |
| YNL191W | DUG3 | Component of glutamine amidotransferase (GATase II); forms a complex | 158 | 171 | 48 | 22 | 0.46 | 1.08 | 0.42 |
| YJR039W | YJR039W | Putative protein of unknown function; the authentic, non-tagged protei | 200 | 209 | 201 | 89 | 0.44 | 1.05 | 0.42 |
| YBR298C | MAL31 | Maltose permease; high-affinity maltose transporter (alpha-glucoside tr | 193 | 354 | 137 | 107 | 0.78 | 1.83 | 0.43 |
| YNL098C | RAS2 | GTP-binding protein; regulates nitrogen starvation response, sporulatior | 124 | 240 | 99 | 84 | 0.85 | 1.94 | 0.44 |
| YER183C | FAU1 | 5,10-methenyltetrahydrofolate synthetase; involved in folic acid biosynt | 206 | 214 | 173 | 79 | 0.46 | 1.04 | 0.44 |
| YNL154C | YCK2 | Palmitoylated plasma membrane-bound casein kinase I isoform; shares | 147 | 140 | 57 | 24 | 0.42 | 0.95 | 0.44 |
| YDR475C | JIP4 | Protein of unknown function; previously annotated as two separate ORF | 184 | 305 | 166 | 123 | 0.74 | 1.66 | 0.45 |
| YPR133W | TOM5 | Component of the TOM (translocase of outer membrane) complex; resp | 166 | 155 | 213 | 89 | 0.42 | 0.93 | 0.45 |
| YBL071C-I | YBL071C-I | Putative protein of unknown function; identified by gene-trapping, micr | 163 | 147 | 265 | 107 | 0.40 | 0.90 | 0.45 |
| YNL230C | ELA1 | Elongin A; F-box protein that forms a heterodimer with Elc1p and is requ | 101 | 195 | 52 | 45 | 0.87 | 1.93 | 0.45 |
| YDL085W | NDE2 | Mitochondrial external NADH dehydrogenase; catalyzes the oxidation of | 166 | 344 | 126 | 118 | 0.94 | 2.07 | 0.45 |
| YBR133C | HSL7 | Protein arginine N-methyltransferase; exhibits septin and Hsl1p-depend | 162 | 130 | 281 | 102 | 0.36 | 0.80 | 0.45 |
| YML071C | COG8 | Component of the conserved oligomeric Golgi complex; a cytosolic tethe | 167 | 125 | 162 | 55 | 0.34 | 0.75 | 0.45 |
| YCR060W | TAH1 | Component of conserved R2TP complex (Rvb1-Rvb2-Tah1-Pih1); R2TP cc | 189 | 191 | 200 | 92 | 0.46 | 1.01 | 0.46 |
| YGR252W | GCN5 | Catalytic subunit of ADA and SAGA histone acetyltransferase complexes; | 206 | 188 | 187 | 78 | 0.42 | 0.91 | 0.46 |
| YGL201C | MCM6 | Protein involved in DNA replication; component of the Mcm2-7 hexamer | 190 | 131 | 269 | 85 | 0.32 | 0.69 | 0.46 |
| YBR172C | SMY2 | Protein of unknown function involved in COPII vesicle formation; interac | 159 | 236 | 166 | 113 | 0.68 | 1.48 | 0.46 |
| YJL179W | PFD1 | Subunit of heterohexameric prefoldin; prefoldin binds cytosolic chapero | 167 | 154 | 81 | 35 | 0.43 | 0.92 | 0.47 |

#### Supplementary Table S2. Yeast Strains.

| Identifier | Genotype | Source |
|---|---|---|
| CTY182 | MATa *ura3-52 lys2-801 his3-Δ200* | (Bankaitis, Malehorn et al., 1989) |
| CTY1-1A | *MATa ura3-52 lys2-801 his3-Δ200 sec14-1ts* | (Bankaitis et al., 1989) |
| PYY19 | *MATa ura3-52 lys2-801 his3-Δ200 sfh4Δ::Ura psd1Δ::KanMX* | This study |
| PYY23 | *MATa ura3-52 lys2-801 his3-Δ200 psd1Δ::KanMX* | This study |
| PYY29 | *MATa ura3-52 lys2-801 his3-Δ200 psd2Δ::KanMX* | This study |
| PYY30 | *MATa ura3-52 lys2-801 his3-Δ200 sfh4Δ::HIS psd1Δ::KanMX* | This study |
| PYY31 | *MATa ura3-52 lys2-801 his3-Δ200 psd1Δ::KanMX scs2Δ::HIS3* | This study |
| PYY32 | *MATa ura3-52 lys2-801 his3-Δ200 psd1Δ::KanMX scs22Δ::HIS3* | This study |
| YWY200 | *MATa ura3-52 lys2-801 his3-Δ200 psd1Δ::KanMX pbi1Δ::HIS3* | This study |
| PYY36 | *MATa ura3-52 lys2-801 his3-Δ200 sfh4Δ::HIS3* | This study |
| PYY40 | *MATa ura3-52 lys2-801 his3-Δ200 sfh4Δ::KanMX* | This study |
| PYY49 | *MATa ura3 met14 lys2 trp1 psd1::TRP1 or psd1Δ-1::TRP1 pstB1* | (Trotter, Wu et al., 1998) |
| PYY52 | *MATa ura3-52 lys2-801 his3-Δ200 psd1Δ::KanMX scs2Δ::HIS3 scs22Δ::NatMX* | This study |
| PYY53 | CTY1-1A *sac1-26 sfh4Δ::KanMX* | This study |

| | | |
|---|---|---|
| PYY54 | PYY49 *sac1Δ::NatMX* | This study |
| PYY55 | PYY30 *sac1Δ::NatMX* | This study |
| PYY84 | *MATa ura3-52 lys2-801 his3-Δ200 sfh4Δ::HIS psd1Δ::KanMX inp51Δ::NatMX* | This study |
| PYY85 | *MATa ura3-52 lys2-801 his3-Δ200 sfh4Δ::HIS psd1Δ::KanMX inp52Δ:: NatMX* | This study |
| PYY86 | *MATa ura3-52 lys2-801 his3-Δ200 sfh4Δ::HIS psd1Δ::KanMX inp53Δ:: NatMX* | This study |
| PYY87 | *MATa ura3-52 lys2-801 his3-Δ200 sfh4Δ::HIS psd1Δ::KanMX inp54Δ:: NatMX* | This study |
| PYY97 | *MATa ura3-52 lys2-801 his3-Δ200 sac1-354::HIS3 psd1Δ::KanMX* | This study |
| YWY34 | *MATa ura3 met14 lys2 trp1 psd1::TRP1 or psd1Δ-1::TRP1 pstB1 scs2Δ::KanMX* | This study |
| YWY35 | *MATa ura3-52 lys2-801 his3-Δ200 sfh4Δ::Ura psd1Δ::KanMX scs2Δ::HIS3* | This study |
| YWY36 | *MATa ura3-52 lys2-801 his3-Δ200 sfh4Δ::Ura psd1Δ::KanMX pbi1Δ::HIS3* | This study |
| YWY37 | *MATa ura3-52 lys2-801 his3-Δ200 psd1Δ::KanMX scs2Δ::His scs22Δ:: NatMX sfh4Δ::URA3* | This study |
| YWY38 | *MATa ura3 met14 lys2 trp1 psd1::TRP1 or psd1Δ-1::TRP1 pstB1 pbi1Δ::URA3* | This study |
| YWY40 | *MATa ura3-52 lys2-801 his3-Δ200 sfh4Δ::HIS psd1Δ::KanMX pbi1Δ::URA3* | This study |
| YWY46 | *MATa ura3-52 lys2-801 his3-Δ200 sfh4Δ::HIS psd1Δ::KanMX scs22Δ::URA3* | This study |
| YWY47 | *MATa ura3 met14 lys2 trp1 psd1::TRP1 or psd1Δ-1::TRP1 pstB1 scs22Δ::URA3* | This study |
| YWY48 | *MATa ura3-52 lys2-801 his3-Δ200 sfh4Δ::Ura psd1Δ::KanMX scs2Δ::HIS3 scs22Δ::NatMX* | This study |

| | | |
|---|---|---|
| YWY49 | *MATa ura3 met14 lys2 trp1 psd1::TRP1 or psd1Δ-1::TRP1 pstB1 scs2Δ::NEO scs22Δ::NatMX* | This study |
| YWY60 | *MATa his3Δ1 leu2Δ0 met15Δ0 ura3Δ0 SFH3-TAP::HIS3MX6* | Dharmacon |
| YWY64 | *MATa ura3-52 lys2-801 his3-Δ200 SFH4-TAP::HIS3* | This study |
| YWY65 | *MATa ura3-52 lys2-801 his3-Δ200* $sfh4^{F175L}$*-TAP::HIS3* | This study |
| YWY66 | *MATa ura3-52 lys2-801 his3-Δ200* $sfh4^{F175A}$*-TAP::HIS3* | This study |
| YWY69 | *MATa ura3-52 lys2-801 his3-Δ200* $stt4^{G1782D}$*::NatMX4* | This study |
| YWY70 | *MATa ura3-52 lys2-801 his3-Δ200 psd1Δ::KanMX* $stt4^{G1782D}$*::NatMX4* | This study |
| YWY71 | *MATa his3Δ1 leu2Δ0 met15Δ0 ura3Δ0 SFH3-TAP::HIS3MX6* $stt4^{G1782D}$*::NatMX4* | This study |
| YWY72 | *MATa ura3-52 lys2-801 his3-Δ200 SFH4-TAP::HIS3MX6* $stt4^{G1782D}$*::NatMX4* | This study |
| YWY73 | *MATa ura3-52 lys2-801 his3-Δ200 sfh4F175L-TAP::HIS3MX6* $stt4^{G1782D}$*::NatMX4* | This study |
| YWY74 | *MATa ura3-52 lys2-801 his3-Δ200 sfh4F175A-TAP::HIS3MX6* $stt4^{G1782D}$*::NatMX4* | This study |
| YWY75 | *MATa ura3-52 lys2-801 his3-Δ200 Δpsd1::KanMX STT4::NatMX4* | This study |
| YWY76 | *MATa ura3-52 lys2-801 his3-Δ200* $sfh4^{T266D,K269A}$*-TAP::HIS3* | This study |
| YWY104 | *MATa ura3-52 lys2-801 his3-Δ200 dpl1Δ::KanMX4* | This study |
| YWY106 | *MATa ura3-52 lys2-801 his3-Δ200 psd1Δ::HIS3* | This study |
| YWY108 | *MATa ura3-52 lys2-801 his3-Δ200 Δect1::KanMX Δpsd1::HIS3* | This study |

| | | |
|---|---|---|
| YWY110 | *MATa ura3-52 lys2-801 his3-Δ200 ect1Δ::KanMX psd2Δ::HIS3* | This study |
| YWY111 | *MATa ura3-52 lys2-801 his3-Δ200 stt4G1782D::NatMX4 psd2Δ::HIS3* | This study |
| YWY112 | *MATa ura3-52 lys2-801 his3-Δ200 psd1Δ::NatMX6* | This study |
| PYY90 | *MATa ura3-52 lys2-801 his3-Δ200 psd1Δ::KanMX osh6Δ::NatMX* | This study |
| YWY128 | *MATa ura3-52 lys2-801 his3-Δ200 sfh4Δ::HIS psd1Δ::KanMX osh6Δ::URA3* | This study |
| YWY129 | *MATa ura3 met14 lys2 trp1 psd1::TRP1 or psd1Δ-1::TRP1 pstB1 osh6Δ::URA3* | This study |
| PYY91 | *MATa ura3-52 lys2-801 his3-Δ200 psd1Δ::KanMX osh7Δ::NatMX* | This study |
| YWY131 | *MATa ura3-52 lys2-801 his3-Δ200 Δsfh4::HIS psd1Δ::KanMX osh7Δ::URA3* | This study |
| YWY132 | *MATa ura3 met14 lys2 trp1 psd1::TRP1 or psd1Δ-1::TRP1 pstB1 osh7Δ::URA3* | This study |
| YWY136 | *MATa ura3-52 lys2-801 his3-Δ200 psd1Δ::KanMX osh1Δ::URA3* | This study |
| YWY137 | *MATa ura3 met14 lys2 trp1 psd1::TRP1 or psd1Δ-1::TRP1 pstB1 osh1Δ::URA3* | This study |
| YWY138 | *MATa ura3-52 lys2-801 his3-Δ200 psd1Δ::KanMX osh2Δ::URA3* | This study |
| YWY139 | *MATa ura3 met14 lys2 trp1 psd1::TRP1 or psd1Δ-1::TRP1 pstB1 osh2Δ::URA3* | This study |
| YWY140 | *MATa ura3-52 lys2-801 his3-Δ200 psd1Δ::KanMX osh3Δ::URA3* | This study |
| YWY141 | *MATa ura3 met14 lys2 trp1 psd1::TRP1 or psd1Δ-1::TRP1 pstB1 osh3Δ::URA3* | This study |
| YWY142 | *MATa ura3-52 lys2-801 his3-Δ200 psd1Δ::KanMX osh4Δ::URA3* | This study |

| | | |
|---|---|---|
| YWY143 | *MATa ura3 met14 lys2 trp1 psd1::TRP1 or psd1Δ-1::TRP1 pstB1 osh4Δ::KanMX4* | This study |
| YWY144 | *MATa ura3-52 lys2-801 his3-Δ200 psd1Δ::KanMX osh5Δ::URA3* | This study |
| YWY145 | *MATa ura3 met14 lys2 trp1 psd1::TRP1 or psd1Δ-1::TRP1 pstB1 osh5Δ::URA3* | This study |
| YWY146 | *MATa ura3-52 lys2-801 his3-Δ200 STT4-TAP::HIS3MX6* | This study |
| YWY147 | *MATa ura3-52 lys2-801 his3-Δ200 psd2Δ::KanMX STT4-TAP::HIS3MX6* | This study |
| YWY148 | *MATa ura3-52 lys2-801 his3-Δ200 sfh4Δ::KanMX STT4-TAP::HIS3MX6* | This study |
| YWY149 | *MATa ura3-52 lys2-801 his3-Δ200* $stt4^{G1782D}$*-TAP::HIS3MX6* | This study |
| YWY152 | *MATa ura3-52 lys2-801 his3-Δ200 sfh4Δ::HIS psd1Δ::KanMX osh1Δ::URA3* | This study |
| YWY153 | *MATa ura3-52 lys2-801 his3-Δ200 sfh4Δ::HIS psd1Δ::KanMX osh2Δ::URA3* | This study |
| YWY154 | *MATa ura3-52 lys2-801 his3-Δ200 sfh4Δ::HIS psd1Δ::KanMX osh3Δ::URA3* | This study |
| YWY155 | *MATa ura3-52 lys2-801 his3-Δ200 sfh4Δ::HIS psd1Δ::KanMX osh4Δ::URA3* | This study |
| YWY156 | *MATa ura3-52 lys2-801 his3-Δ200 sfh4Δ::HIS psd1Δ::KanMX osh5Δ::URA3* | This study |
| SEY6210.1 | *MATa leu2-3,112 ura3-52 his3-Δ200 trp1-Δ901 lys2-801 suc2-Δ9* | (Stefan, Audhya et al., 2002) |
| ANDY198 | *SEY6210.1 ist2Δ::HISMX6 scs2Δ::TRP1 scs22Δ::HISMX6 tcb1Δ::KANMX6 tcb2Δ::KANMX6 tcb3Δ::HISMX6* | (Manford, Stefan et al., 2012) |
| YWY167 | *SEY6210.1 psd1Δ::NatMX6* | This study |
| YWY168 | *ANDY198 psd1Δ::NatMX6* | This study |

| | | |
|---|---|---|
| YWY169 | *SEY6210.1 psd2Δ::NatMX* | This study |
| YWY170 | *ANDY198 psd2Δ::NatMX* | This study |
| YWY182 | *MATa ura3-52 lys2-801 his3-Δ200 psd1Δ::HIS3 vps13Δ::KanMX4* | This study |
| YWY193 | *MATa ura3-52 lys2-801 his3-Δ200 psd1Δ::HIS3 lam5Δ::KanMX4* | This study |
| YWY197 | *MATa ura3-52 lys2-801 his3-Δ200 psd1Δ::HIS3 lam6Δ::KanMX4* | This study |

#### Supplementary Table 3. Plasmid List.

| Identifier | *Recombinant DNA* | Source |
|---|---|---|
| pYW18 | *pET28b SFH4* | This study |
| pPY126 | *pET28b* $sfh4^{T266D\ K269A}$ | This study |
| pPY127 | *pET28b* $sfh4^{T266W\ K269A}$ | This study |
| pPY130 | *pRS316 STT4* | This study |
| pPY131 | *pRS316* $stt4^{G1782D}$ | This study |
| pPY132 | *pRS316* $stt4^{D1754A}$ | This study |
| pPY195 | *pET28b* $sfh4^{F175L}$ | This study |
| pYW13 | *YEp352 PSD2-HA* | (Gulshan, Shahi et al., 2010) |
| pYW31 | *pRS316 SFH4* | This study |
| pYW32 | *pRSII426 SFH4* | This study |
| pYW35 | *pRS316* $SFH4^{F175L}$ | This study |
| pYW36 | *pRS316* $SFH4^{F175A}$ | This study |
| pYW37 | *pRS316* $SFH4^{F175I}$ | This study |
| pYW38 | *pRS316* $SFH4^{F175R}$ | This study |
| pYW39 | *pRS316* $SFH4^{F175E}$ | This study |
| pYW40 | *pRS316* $SFH4^{F175W}$ | This study |
| pYW41 | *pRS316* $SFH4^{F175H}$ | This study |
| pYW42 | *pRS316* $SFH4^{F175Y}$ | This study |
| pYW43 | *pRSII426* $SFH4^{F175L}$ | This study |
| pYW44 | *pRSII426* $SFH4^{F175A}$ | This study |
| pYW46 | *pRSII426* $SFH4^{F175R}$ | This study |
| pYW47 | *pRSII426* $SFH4^{F175E}$ | This study |
| pYW58 | *pRS316* $PGK1_{Pro}$-$SFH4^{F175L}$ | This study |

| | | |
|---|---|---|
| pYW59 | *pRS316 PGK1$_{Pro}$-SFH4$^{F175A}$* | This study |
| pYW61 | *pRS316 PGK1$_{Pro}$-SFH4$^{F175R}$* | This study |
| pYW62 | *pRS316 PGK1$_{Pro}$-SFH4$^{F175E}$* | This study |
| pYW66 | *pET28b sfh4$^{F175A}$* | This study |
| pYW67 | *pET28b sfh4$^{F175R}$* | This study |
| pYW68 | *pET28b sfh4$^{F175E}$* | This study |
| pYW85 | *pBluescript II SK(+) SFH4-TAP::HIS3* | This study |
| pYW86 | *pBluescript II SK(+) SFH4-TAP::HIS3* | This study |
| pYW88 | *pBluescript II SK(+) sfh4$^{F175L}$-TAP::HIS3* | This study |
| pYW89 | *pBluescript II SK(+) sfh4$^{F175A}$-TAP::HIS3* | This study |
| pYW90 | *pBluescript II SK(+) sfh4$^{F175R}$-TAP::HIS3* | This study |
| pYW91 | *pBluescript II SK(+) sfh4$^{F175E}$-TAP::HIS3* | This study |
| pYW93 | *pBluescript II SK(+) stt4G1782D::NatMX* | This study |
| pYW97 | *pRS316-sfh4$^{T266D,K269A}$* | This study |
| pYW109 | *pBluescript II SK(+) ~ sfh4$^{T266D,K269A}$-TAP::HIS3* | This study |
| pYW127 | *pBluescript II SK(+) ~ STT4-TAP::HIS3* | This study |
| pYW128 | *pBluescript II SK(+) ~ stt4$^{G1782D}$-TAP::HIS3* | This study |
| pYW134 | *YCplac111 PACT1-GFP-Myc-HMH-RitC* | (Quon, Sere et al., 2018) |
| pYW142 | *pRSII426 YPP1* | This study |
| pYW143 | *pRSII426 EFR3* | This study |
| pYW144 | *pRSII426 SFK1* | This study |
| pYW145 | *pDR195 OSH6* | This study |
| pYW146 | *pDR195 OSH7* | This study |
